# High-fat circulating nutrients promote growth and invasion in a 3D microfluidic tumor model of triple-negative breast cancer

**DOI:** 10.1101/2025.07.10.664224

**Authors:** Maryam Kohram, Carolina Trenado-Yuste, Molly C. Brennan-Smith, Evelyn S. Navarro Salazar, Pengfei Zhang, Jasmine E. Hao, Xincheng Xu, Bharvi Chavre, William Oh, Sherry X. Zhang, Susan E. Leggett, Rolf-Peter Ryseck, Joshua D. Rabinowitz, Celeste M. Nelson

## Abstract

Diet influences the levels of small molecules that circulate in plasma and interstitial fluid, altering the biochemical composition of the tumor microenvironment (TME). These circulating nutrients have been associated with how tumors grow and respond to treatment, but it remains difficult to parse their direct effects on cancer cells. Here, we combine a three-dimensional (3D) microfluidic tumor model with physiologically relevant culture media to investigate how concentrations of circulating nutrients influence tumor growth, cancer cell invasion, and overall tumor metabolism. Human triple-negative breast cancer cells cultured in 2D under media conditions mimicking five different dietary states show no observable differences in proliferation or morphology. Nonetheless, those exposed to high-fat conditions exhibit increased metabolic activity and upregulate genes associated with motility and extracellular matrix remodeling. In the 3D microfluidic model, high-fat conditions accelerate tumor growth and invasion and induce the formation of hollow cavities. Surprisingly, the presence of these cavities does not correlate with an increase in apoptosis or ferroptosis. Instead, RNA-sequencing analysis revealed that high-fat conditions induce the expression of *MMP1*, consistent with cavitation via cell invasion.

Mimicking the flow of circulating nutrients within the TME can thus be used to identify novel connections between metabolic states and tumor phenotype.

## Introduction

Tumor progression is often slowed by targeting cancer cells directly or indirectly using chemotherapy or radiation. The response of a tumor to these treatments depends in part on the surrounding microenvironment. Dietary composition affects the levels of nutrients and metabolites that circulate in plasma and interstitial fluid and, thus, the availability of small molecules to the cells within the tumor^1,2^. The presence of metabolites in the tumor microenvironment (TME) is likely to affect the progression and responsiveness of cancer cells to different treatments^3,4^. Therefore, understanding the effects of circulating nutrients on tumor cell behavior could suggest new means of slowing disease progression^5–8^.

Unfortunately, disentangling the relative effects of circulating nutrients on tumor progression in patients and mice is challenging. Clinical trials often overlook individual factors such as diet^9^, with limited studies focused on insulin-related dietary factors or changes in carbohydrate intake after breast cancer diagnosis^10^. A larger number of studies have been conducted using mouse models of cancer and have suggested a potential for diets to impact overall survival^11–15^.

Specifically, high-fat diets have been found to accelerate breast cancer progression in tumor-bearing mice^16^, regardless of obesity status^17^. In contrast, ketogenic diets have been found to reduce the growth of breast tumors^18^, but paradoxically enhance the metastatic capacity of breast cancer cells^19^. The effectiveness of a ketogenic diet may vary by mouse strain and sex^20^, highlighting the need for a more detailed understanding of the synergy between diet and tumor progression. Diet also affects other features of the TME, including the immune system, microbiome, and adipose tissue, all of which likely contribute to the interpretation of its effects on tumor progression. Because of this complexity, isolating the direct impact of circulating nutrients and metabolites on tumor cells remains challenging.

The TME is characterized by a continuous flow of interstitial fluid that transports small molecules to and from cancer cells^21,22^. This convective transport adjusts the availability and concentration of nutrients, impacting tumor behavior and responsiveness to treatments.

Accurately replicating the flow of fluid within the TME is therefore essential for modeling tumor responses to dietary interventions. In mouse models, it is challenging to alter individual components within a diet and to track the phenotype of an individual tumor as it grows^8,23^. In contrast, culture models allow for one to examine the effects of dietary interventions on human cancer cells over time, with higher throughput than animal models^24^. However, culture models also present their own limitations. Standard culture methods, such as spheroids embedded in collagen, rely upon the intermittent delivery of nutrients. As a result, nutrient concentration in these models is neither consistently maintained nor accurately controlled, failing to replicate the continuous, circulating nutrient flow observed *in vivo*. As such, accurately studying dietary effects in culture requires a tumor model which replicates the continuous flow of nutrients within the interstitial fluid.

Here, we took advantage of human plasma-like media^25^ and mimicked the circulating nutrients characteristic of five different dietary conditions: baseline, post-prandial, diabetic, ketogenic, and high-fat. Under 2D culture, MDA-MB-231 human triple-negative breast cancer (TNBC) cells exhibited similar morphologies and proliferation rates across the media conditions. However, RNA-sequencing analysis revealed differences in gene expression, with high-fat conditions upregulating genes associated with motility and extracellular matrix (ECM) remodeling. These transcriptional changes were accompanied by elevated metabolic activity in high-fat media, despite mass spectrometry analysis showing largely similar profiles of secreted metabolites across all conditions. We next mimicked the continuous delivery of circulating nutrients by culturing cells as three-dimensional (3D) aggregates in a microfluidic system that generated interstitial fluid flow comparable to that observed *in vivo*^26–33^. Under high-fat conditions, tumors exhibited accelerated growth and invasion, without appreciable differences in proliferation compared to baseline conditions. Notably, high-fat conditions also led to the formation of hollow cavities within the tumors. These structures did not correlate with increased rates of apoptosis or ferroptosis but were instead associated with elevated expression of *MMP1*, which we hypothesize promoted ECM degradation and subsequent cancer cell dispersal into the surrounding microenvironment.

## Materials and Methods

### Cell culture

MDA-MB-231 human breast cancer cells (ATCC) were maintained in a humidified incubator at 37℃ and 5% CO_2_ in one of six different media conditions: DMEM, baseline, post-prandial, diabetic, ketogenic, or high fat. Specifically, cells were cultured in either DMEM/F12 medium (Hyclone) supplemented with 10% fetal bovine serum (FBS, R&D Systems) and 1% gentamicin (Gibco), or human plasma-like medium (HPLM; **Table S1**)^25^ supplemented with 10% FBS dialyzed by ultrafiltration (Sigma-Aldrich), 1% gentamicin (Gibco), and the following additives: 5-ng/mL bovine pancreas insulin (Sigma-Aldrich; post-prandial); 10-mM D-glucose (Sigma-Aldrich; diabetic); 2-mM DL-β-hydroxybutyric acid sodium salt (BHB; Sigma-Aldrich; ketogenic); or 4% (w/v) lipid-rich bovine serum albumin (BSA, AlbuMAX^TM^ I, Gibco; high fat, **Table S2**). Cells were passaged at a 1:3-6 ratio and used before passage 25. MDA-MB-231 cells cultured in baseline HPLM were tested for authentication by short tandem repeat (STR) profiling using the ATCC Cell Line Authentication Service. The STR profiles were compared to the reference profiles provided by ATCC, confirming the identity of the cell line.

### Viscosity measurements

The viscosity of each media formulation was measured at room temperature using an Ostwald viscometer (Ace Glass, Inc.). Viscosity (𝜂) was calculated using the equation 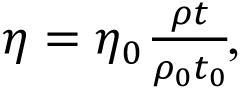 where 𝜂_0_ = 1 cP is the viscosity of water at room temperature, 𝜌 and 𝜌_0_are the densities of the medium and water, respectively, and 𝑡 and 𝑡_0_are the times of outflow of the medium and water, respectively. The measurements were performed three times using three independently prepared media samples. The uncertainty for each measurement was determined using the propagation of errors using the equation 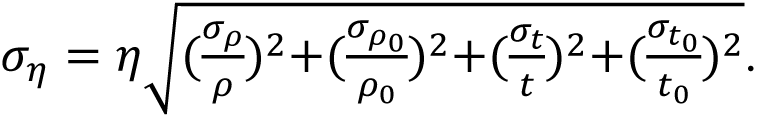

### Microfluidic tumor model

Microtumors comprised of MDA-MB-231 cells were generated using a microfluidic approach, as described previously^28,30,31^. Briefly, a silicone mold was used to create channels within polydimethylsiloxane (PDMS), which were exposed to UV/ozone and bonded to a glass-bottomed tissue-culture dish. The channels were then immediately coated with 1-mg/mL poly-D-lysine hydrobromide (Sigma-Aldrich). Acupuncture needles (Austin Medical; 120 𝜇m in diameter) were coated with 1% BSA in phosphate buffered saline (PBS) and positioned within one of the PDMS channels. A 4-mg/mL solution of type I collagen (Koken) was prepared using 10-N NaOH, 10X Hank’s Balanced Salt Solution (HBSS; Gibco), MilliQ water, and HPLM, resulting in an aqueous solution of collagen with a pH of 9.5. The collagen solution was introduced into the PDMS channels and incubated at 37°C for 20 minutes to form a gel. The needles were subsequently removed to yield two empty cavities, one on each side of the gel. A suspension of MDA-MB-231 cells in baseline culture medium supplemented with 100-µg/mL DNase I (Roche) was added to fill the cavity on one side of the gel, and then the excess cells were washed away. Samples were cultured for 48 hours, after which the flow of interstitial fluid was initiated, as previously described^28,30,31^. Tumors were monitored for invasions and conditioned medium was collected every 12 hours on days 3.5-5.5 from the downstream well for mass spectrometry analysis, while fresh medium was added to the upstream well.

The permeability of the collagen gel was calculated using Darcy’s Law 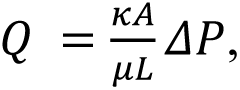 where 𝑄 is the volumetric flow rate of the culture medium, 𝜅 is the permeability of the collagen gel, 𝐴 is the cross-sectional area of the collagen gel, 𝜇 is the viscosity of the culture medium, 𝐿 is the length of the gel, and 𝛥𝑃 is the pressure difference across the gel. This approach revealed that the permeability of the collagen gel was 0.100 ± 0.0076 µm^2^ resulting in a flow rate of 5 – 8 𝜇L/hr, which is on the same order of magnitude as previously reported values^28,29^.

### Immunofluorescence analysis

To assess morphology and phenotype in 2D culture, cells were seeded at 4,000 cells per well in a 96-well plate and cultured for 72 hours under each of the five different media conditions.

Samples were fixed for 15 minutes in 4% paraformaldehyde (PFA) in PBS and then washed three times with PBS. Fixed samples were incubated for 1 hour at room temperature in blocking buffer comprised of 20% BSA and 10% normal goal serum (NGS) in PBS supplemented with 0.3% Triton X-100 (PBST). Blocked samples were washed with PBST and incubated overnight with primary antibodies against E-cadherin (Cell Signaling, 24E10 rabbit mAb #3195) and vimentin (Sigma-Aldrich, clone LN-6 mouse V2258) diluted 1:200 in blocking buffer, followed by additional washing with PBST. Samples were incubated in the dark for 1 hour at room temperature with Alexa Fluor-conjugated secondary antibodies and phalloidin (Thermo Fisher Scientific) diluted 1:200 in blocking buffer. Samples were washed twice with PBS, counterstained with Hoechst 33342 (Invitrogen) diluted 1:1000 in PBS for 15 minutes, and then washed twice with PBS.

Proliferation assays were conducted using the Click-iT EdU Alexa Fluor 555 Kit (Thermo Fisher Scientific). Samples were exposed to EdU for 2 hours and then immediately fixed with 4% PFA in PBS before applying the Click-iT cocktail reagent. 2D samples were then counterstained with Hoechst 33342 diluted 1:1000 in PBS. Engineered tumor samples were incubated overnight at 4°C with Hoechst 33342 diluted 1:1000 in 1% BSA in PBST. Subsequently, the tumor samples were incubated for 30 minutes in 25% glycerol in PBS, and then 50% glycerol in PBS, and then transferred into 75% glycerol in PBS and stored at 4°C for future imaging.

Lipid peroxidation was assessed using the Click-iT Lipid Peroxidation Imaging Kit – Alexa Fluor 488 (Thermo Fisher Scientific). Tumors were exposed to linoleamide alkyne for 2 hours, washed three times with PBS, fixed with 4% PFA in PBS, and then washed three times with PBS. Samples were incubated in 0.1% PBST for one hour, blocked with 1% BSA in PBS for one hour, and then washed twice with PBS. Subsequently, tumors were incubated in Click-iT reaction cocktail for one hour and washed twice with 1% BSA in PBS and twice with PBS. To label apoptotic cells, tumors were first incubated in blocking buffer comprised of 10% NGS in 0.1% PBST overnight at 4°C. Blocked tumors were incubated overnight with primary antibody against cleaved caspase-3 (Cell Signaling, Asp175 rabbit 9661) diluted 1:400 in blocking buffer. The following day, samples were washed four times with 0.1% PBST and incubated overnight at 4°C in Alexa Fluor 647 conjugated secondary antibody (Invitrogen, goat anti-rabbit A21245) diluted 1:200 in blocking buffer. Samples were washed four times with 0.1% PBST and counterstained with Hoechst 33342 diluted 1:2000 in PBS for 15 minutes, washed once with PBS, and transferred into 75% glycerol in PBS using the same procedure as described above.

### Analysis of energy metabolism

The oxygen consumption rate (OCR) and extracellular acidification rate (ECAR) of cells in the different media conditions were measured using a Seahorse XFe24 Analyzer (Agilent). The Seahorse XF Glucose/Pyruvate Oxidation Stress Test Kit (Agilent) was used according to the manufacturer’s protocol. Cells were seeded at a density of 100,000 cells/well in a Seahorse XF24 V28 PS Cell Culture Microplate (Agilent) in their corresponding media conditions one day before measurement. One hour before the measurements, the media was replaced with HPLM without phenol red and bicarbonate, adjusted to a pH of 7.4, for all five media conditions. For the cells cultured in DMEM, the media was replaced with DMEM (Agilent) supplemented with 10-mM glucose, 1-mM pyruvate, and 2-mM glutamine. The plates were incubated at 37°C for one hour prior to measurements. OCR and ECAR were measured every 8 minutes during basal respiration and after sequential injection of the relevant pathway inhibitors: 2-𝜇M UK5099, 1.5-𝜇M oligomycin, 1.0-𝜇M carbonyl cyanide-4 (trifluoromethoxy) phenylhydrazone (FCCP), and 0.5-𝜇M rotenone/antimycin A.

### Bulk RNA-sequencing analysis

Cells were cultured in six different media conditions in a 6-well tissue-culture plate at a density of 200,000 cells/well. After 72 hours, RNA was extracted and purified using an RNeasy Mini Kit (Qiagen) according to the manufacturer’s protocol. The extracted RNA was processed by the Princeton University Genomics Core Facility. RNA quality was evaluated using an Agilent Bioanalyzer. RNA-Seq directional library preparation was performed on an Apollo 324 Robot and the samples were run in a single lane on a NovaSeq SP 100 nucleotide Flowcell v1.5 with a read depth of ∼650-800 million reads.

Reads were aligned to the human reference genome (GRCh38.p14) using STAR, followed by transcriptome quantification using HTSeq-Count. Raw read counts were normalized using the estimateSizeFactors function in the DESeq2 package^34^. Normalized counts were transformed into regularized log for variance stabilization before performing principal component analysis (PCA) and Z-score heatmaps. Differentially expressed genes (DEGs) were identified using Wald tests, followed by shrinkage of log fold-change using the apeglm method to avoid inflation of fold changes of genes with low counts^34,35^. Significant DEGs had a Benjamini-Hochberg-adjusted p-value <0.05 and an absolute log2 fold change >2. Over-representation analysis (ORA) in Gene Ontology (GO) terms of biological processes was conducted separately for the downregulated and upregulated significant DEGs of the high-fat condition in reference to the baseline condition using the gprofiler2 package^36^.

### Sectioning tumors

Tumor samples were fixed, placed in a disposable cryomold, and fully surrounded with Optimal Cutting Temperature (OCT) compound to ensure complete coverage. Cryomolds containing tumors were rapidly frozen using dry ice. Once frozen, cryomolds were stored at-80°C until sectioning. Frozen tumors were cut into ∼50-µm-thick sections using a CM3050S cryostat. After sectioning, tumor sections were placed onto Fisherbrand Superfrost Plus microscope slides (Fisher Scientific) and stored at-80°C until use.

### Hematoxylin and eosin (H&E) staining

H&E staining was performed using a modified version of a previously described protocol^37^. Slides with tumor sections were placed in staining racks at room temperature for 30 minutes. Tumor sections were fixed again in 4% PFA in PBS for 10 minutes. Fixed sections were washed twice with PBS and stained in Harris Hematoxylin solution for 2 minutes. Stained samples were rinsed with 1-2 dips in acid alcohol. Sections were washed three times in Milli-Q water, stained with bluing solution, and washed three times in Milli-Q water. Sections were dehydrated by sequential immersion in 75%, 80%, and 95% alcohol. Dehydrated sections were stained with Eosin Y for 20 seconds, and then washed by immersing once in 95% alcohol, then three times in absolute alcohol. Sections were cleared in two successive xylene substitute (Electron Microscopy Sciences) baths. Finally, sections were mounted under a coverslip with acrylic mounting media (Sigma-Aldrich).

### Prussian blue staining

For Prussian blue staining, slides with tumor sections were placed in staining racks at room temperature for 30 minutes. Tumor sections were fixed again in 4% PFA in PBS for 10 minutes, and then washed twice with PBS. Sections were stained for 3 minutes using a working iron-stain solution prepared by mixing equal volumes of potassium ferrocyanide and hydrochloric acid (Iron Stain Kit, Prussian Blue stain ab150674). Sections were then washed three times in Milli-Q water, followed by staining with nuclear fast red. Samples were then washed and dehydrated by sequential immersion in 75%, 80%, and 95% alcohol. Finally, sections were mounted under a coverslip with acrylic mounting media.

### Fluorescence in situ hybridization

Fluorescence in situ hybridization was performed using the standard RNAscope Multiplex Fluorescent V2 Assay (Advanced Cell Diagnostics) protocol for fixed-frozen samples. The probe was for *Homo sapiens MMP1* (channel 1, 1057851) and the fluorophore was Opal 650 (Akoya Biosciences, FP1496001KT).

### Imaging

Engineered tumors were imaged daily using a Nikon Ti-U microscope equipped with a digital CMOS camera (Hamamatsu ORCA-Flash 4.0 V3) using a 10× air objective. Fluorescent samples were imaged using a Nikon Eclipse Ti2 microscope equipped with a spinning disk confocal (X-light V2, Crest Optics) and an ORCA charge-coupled device camera (Hamamatsu, Japan). Images of 2D samples were acquired using either a Plan Fluor 20× 0.45 *NA* or Plan Fluor 40× 1.3 *NA* oil objective. Confocal stacks of tumors (200-µm-deep) were acquired using a Plan Fluor 20× 0.95 *NA* water objective with a step size of 2 µm. Tumors stained for cleaved-caspase-3 and lipid peroxidation were imaged on a Nikon Ti-E microscope attached to a Maico MEMS confocal unit (Hamamatsu, Japan) using a Fluor 40× 1.3 *NA* oil objective.

H&E and Prussian blue-stained sections were imaged using a Leica M205 stereoscope equipped with a MC170 HD camera (25195014 DOC2) using a Leica PLANAPO 2.0× objective (M-series). RNAscope sections were imaged on a Nikon Ti-E microscope attached to a Maico MEMS confocal unit using a Fluor 40× 1.3 *NA* oil objective.

### Image analysis

To quantify proliferation of cells cultured in 2D, the number of EdU-positive cells and total number of nuclei were counted in ImageJ using the StarDist plugin. The percentage of EdU-positive cells was given by the ratio of EdU-positive cells to the total number of nuclei for all images of each condition in each replicate. To quantify proliferation of cells in the 3D engineered tumors, confocal stacks were imported into Imaris. The number of EdU-positive cells and total number of nuclei were counted using the Spots function, with subsequent analysis limited to the initial 200 µm of the tumor and anything extending beyond that point along its axial length.

Tumor morphology was analyzed using a custom image-analysis pipeline^27^. Briefly, a cutoff point for each tumor was delineated as 200 µm into the tumor in the direction of flow at the time of seeding. The total tumor and tumor core that extended beyond this cutoff point were segmented from images acquired on day 4 of culture. The tumor core was defined as the dense region of the tumor excluding any invasions. The area of invasions was calculated as the area of the total tumor remaining after the tumor core was dilated by 25 µm to account for small invadopodia extending from the tumor core and then subtracted from the area of the total tumor.

### Glucose uptake assay

MDA-MB-231 cells were seeded at 200,000 cells/well in a 6-well-plate. After 72 hours, media was collected and stored at-20°C. Media collected from cell-free wells were used as a control and stored at-20°C. Glucose uptake was measured using the Glucose Assay Kit (Abcam ab65333) according to the manufacturer’s instructions.

### Mass spectrometry

Conditioned medium was harvested from the downstream well of the microfluidic device beginning on day 3.5 after seeding and continuing through day 5.5. Control medium was harvested by adding 150 𝜇L of media into a 96-well plate at the same time as fresh media was added to the microtumors. This plate was then placed in the same incubator as the tumors, and the media was collected at the same time as the conditioned media was collected from the tumors. Media collected from tumors in each condition were combined and stored at 4°C until the end of the experiment. The samples were then preserved at-20°C. On the day prior to mass spectrometry, samples were thawed, diluted 1:29 in methanol (Fisher Chemical), and stored at - 80°C overnight. The next day, samples were centrifuged at 19,930g for 20 min at 4°C and the supernatant was collected.

Metabolites were separated by hydrophilic interaction liquid chromatography (HILIC) with an XBridge BEH amide column (2.1-mm × 150-mm, 2.5-𝜇m particle size; Waters, 186006724). The column temperature was set at 25°C. Solvent A was 95:5 (v:v) H_2_O:acetonitrile (with 20-mM ammonium acetate, 20-mM ammonium hydroxide, pH 9.4). Solvent B was acetonitrile. Flow rate was 0.15 mL/min. The LC gradient was: 0-2 min, 90% B; 3-7 min, 75% B; 8-9 min, 70% B; 10-12 min, 50% B; 12-14 min, 25% B; 16-20.5 min, 0.5% B; 21-25 min, 90%. MS analysis was performed on an Orbitrap Exploris 480 mass spectrometer. Full scan was performed in negative mode, at the m/z of 70-1000. The automatic gain control (AGC) target was 1000% and the maximum injection time was 500 ms. The orbitrap resolution was 180,000. Raw mass spectrometry data were converted to mzXML format by MSConvert (ProteoWizard). Pick-peaking was done on El Maven (v0.8.0, Elucidata). Peak-intensity values for the conditioned media corresponding to each media composition were normalized to their respective unconditioned control media using the equation 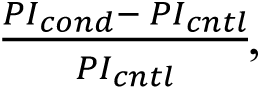 where 𝑃𝐼𝑐𝑜𝑛𝑑 is the peak intensity of the conditioned media, and 𝑃𝐼_𝑐𝑛𝑡𝑙_ is the peak intensity of the control media.

## Statistical analysis

All statistical analyses were performed using GraphPad Prism. The metrics for 2D and 3D morphology, OCR and ECAR values, the percentage of EdU-positive cells, total number of nuclei, percentage of cleaved-caspase-3 positive and percentage of lipid peroxidation in 3D were compared using one-way ANOVA with Tukey’s multiple comparison test. When the result of Bartlett’s test was significant (p<0.05), the Brown-Forsythe ANOVA test was performed and followed by Dunnett’s T3 multiple comparisons test. Otherwise, the one-way ANOVA test and Tukey’s multiple comparisons test were performed. When comparing the maximum distances of invasion, the Bartlett’s test and Brown-Forsythe ANOVA test could not be performed, so the one-way ANOVA test and Tukey’s multiple comparisons test were used. ATP production rates were compared using two-way ANOVA. The rates of invasion were compared using the log-rank (Mantel-Cox) test. Significance for all comparisons was considered p<0.05, and p-values were adjusted for multiple comparisons.

## Results

### MDA-MB-231 cells show similar morphologies and proliferation rates when cultured in 2D in media mimicking different dietary states

Dysregulated cellular metabolism is one of the hallmarks of cancer^38^. Metabolism can be influenced by the circulating nutrients and metabolites available in the tissue microenvironment^8^, which are controlled in part through diet. To study the effects of circulating nutrients on cancer cell behavior, we used media that mimicked the plasma composition of patients with different dietary states. Specifically, we mimicked baseline plasma by taking advantage of human plasma-like medium (HPLM), which is formulated to contain polar metabolites and salts at concentrations observed in human plasma (**Table S1**)^25^. We mimicked post-prandial plasma and the plasma of diabetic patients by adding insulin and glucose. We also mimicked the plasma of patients on ketogenic and high-fat diets by supplementing HPLM with ketones and fatty acids.

Breast cancer motility increases in response to elevated extracellular fluid viscosity, which promotes actin remodeling at the leading edge of the cell^39^. We therefore began by measuring the viscosities of the five different media formulations to ensure that any observed effects on cell morphology could be attributed to differences in biochemical composition rather than differences in viscosity. We found the viscosity of the baseline condition at room temperature to be 1.06 ± 0.02 cP, which is approximately 3.2% higher than that of DMEM supplemented with FBS (**Fig. S1A**). We also found that the high-fat condition had a slightly higher viscosity (∼17.5%) than baseline, consistent with *in vivo* observations, as high-fat diets are known to increase the viscosity of plasma in patients^40^. Nonetheless, the viscosities of all the media conditions were below the thresholds previously found to alter cell motility (>3cP at 37°C)^39^.

We therefore proceeded to establish cultures of MDA-MB-231 human breast cancer cells in each of the different media formulations (**Fig. 1A**). To determine whether the media conditions altered cellular morphology in 2D, we measured projected cellular and nuclear areas and circularity.

**Figure 1.**
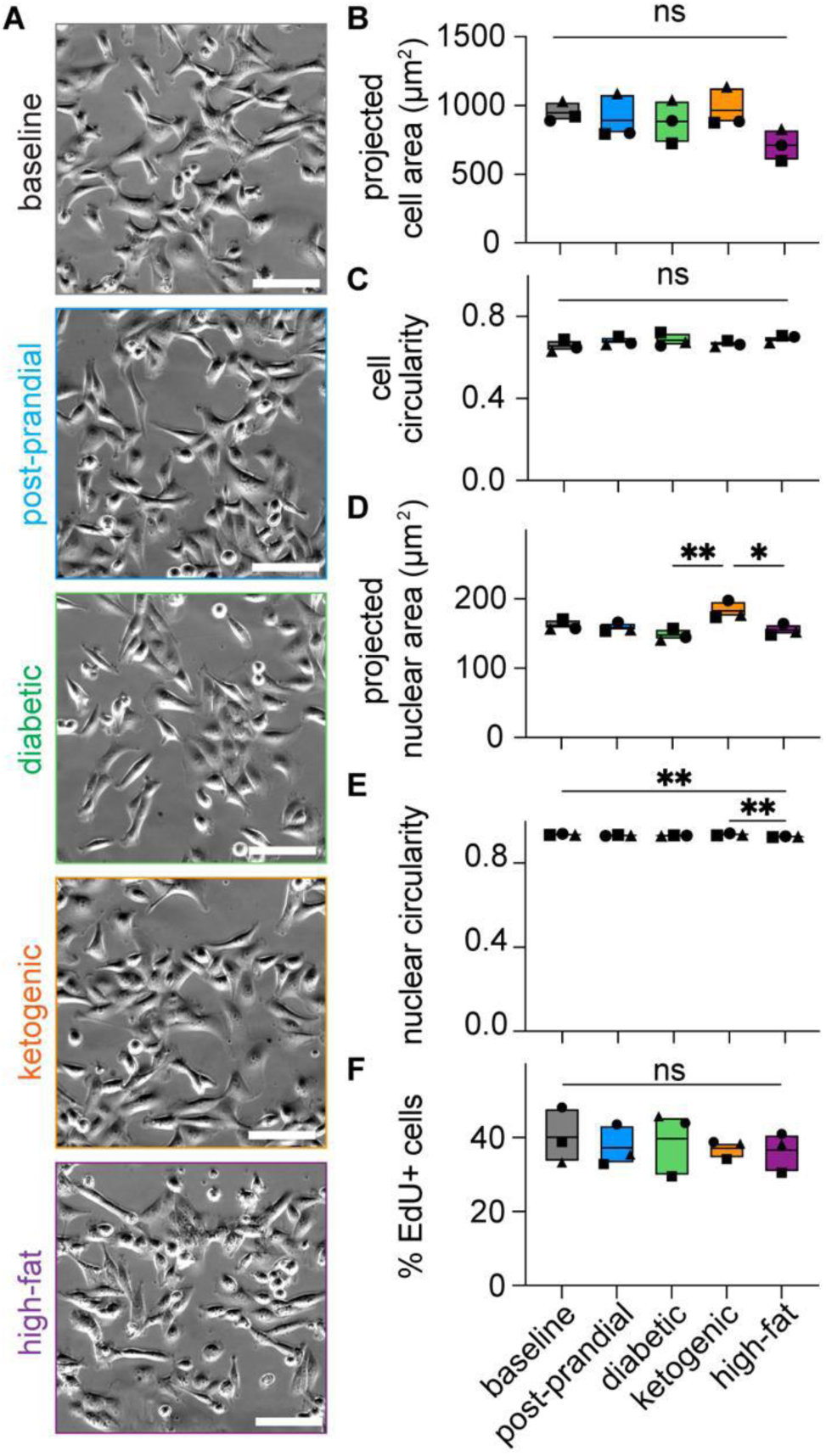
MDA-MB-231 human breast carcinoma cells show similar morphologies and rates of proliferation in media that mimic different circulating nutrient conditions. (**A**) Phase-contrast images of MDA-MB-231 cells cultured in 2D under different media conditions; scale bars = 100 µm. Graphs of (**B**) projected cell area, (**C**) cell circularity, (**D**) nuclear area, and (**E**) nuclear circularity for MDA-MB-231 cells cultured in 2D under different media conditions, pooled over n = 3 independent experiments. Shown are mean and range. (**F**) Graph showing percentage of EdU^+^ MDA-MB-231 cells cultured in 2D under different media conditions, pooled over n = 3 independent experiments. Shown are mean and range. (*), p<0.05; (**), p<0.01; ns, not significant.

This analysis revealed that the projected cell area and circularity remained consistent across all media conditions (**Fig. 1B, C**), with cells displaying a spindle-like morphology (**Fig. 1A**). Cells in the ketogenic media exhibited a slight increase in nuclear area (**Fig. 1D**) while those in the high-fat media exhibited a slight decrease in nuclear circularity (**Fig. 1E**), as compared to baseline. Consistent with the spindle-like morphology, immunofluorescence analysis revealed that cells in all conditions maintained expression of the mesenchymal marker, vimentin (**Fig. S1B**)^41^. To determine whether the media conditions differentially affected cell proliferation, we carried out EdU analysis. This analysis revealed a uniform rate of proliferation, with ∼40% of cells incorporating EdU over a two-hour pulse, irrespective of the media in which the cells were cultured (**Fig. 1F**). These findings suggest that the overall mesenchymal phenotype of MDA-MB-231 cells remains stable across the different media formulations.

### Culture in media mimicking different dietary states alters gene expression in MDA-MB-231 cells

Since the five media conditions resulted in no appreciable differences in cell morphology or proliferation, we turned to RNA-sequencing as an unbiased approach to uncover possible differences in gene expression. This analysis revealed significant transcriptional changes upon transitioning cells from DMEM to HPLM (**Fig. S2A**). Directly comparing the baseline HPLM condition with DMEM revealed 1139 genes that were differentially expressed, including 292 genes upregulated and 847 genes downregulated by culture in HPLM (**Fig. S2B, C**). This shift in gene expression suggests that cells undergo metabolic, structural, and signaling adaptations when transitioned to the more physiologically relevant biochemical microenvironment (**Fig. S3A; Supplementary Text**).

Our RNA-sequencing analysis also revealed differences in gene expression between cells cultured in media mimicking the different dietary conditions, despite the fact that we observed no differences in morphology or proliferation. Specifically, principal component analysis (PCA) of the 1,000 most variable genes revealed that cells cultured under the high-fat condition displayed a transcriptional signature distinct from those under the other conditions (**Fig. 2A**). A heatmap of differentially expressed genes further highlighted gene-expression changes induced by the high-fat media (**Fig. 2B**). To assess condition-specific transcriptional changes, we separately compared each media formulation to baseline HPLM (**Fig. 2C, Fig. S3B-D; Supplementary Text**). Consistent with our PCA analysis, we found that the high-fat condition resulted in the most pronounced transcriptional changes relative to baseline (**Fig. 2C**). Cells cultured under high-fat conditions altered the expression of several genes associated with fatty acids and lipid metabolism (**Fig. 2C**). Specifically, we observed upregulation of *PDK4*, which encodes pyruvate dehydrogenase kinase 4, an enzyme that phosphorylates and inactivates the pyruvate dehydrogenase complex, thereby shifting cellular metabolism from glucose oxidation toward fatty acid utilization^42^. Similarly, we observed increased expression of *ANGPTL4*, which encodes a secreted glycoprotein induced by fatty acids that inhibits lipoprotein lipase, regulating lipid metabolism and glucose homeostasis^43^. Conversely, high-fat conditions led to the downregulation of *SCD* and *PCSK9*, both of which are involved in lipid metabolism, which is consistent with the abundance of lipids in this media condition, potentially reducing the need for *de novo* lipid synthesis and cholesterol regulation.

**Figure 2.**
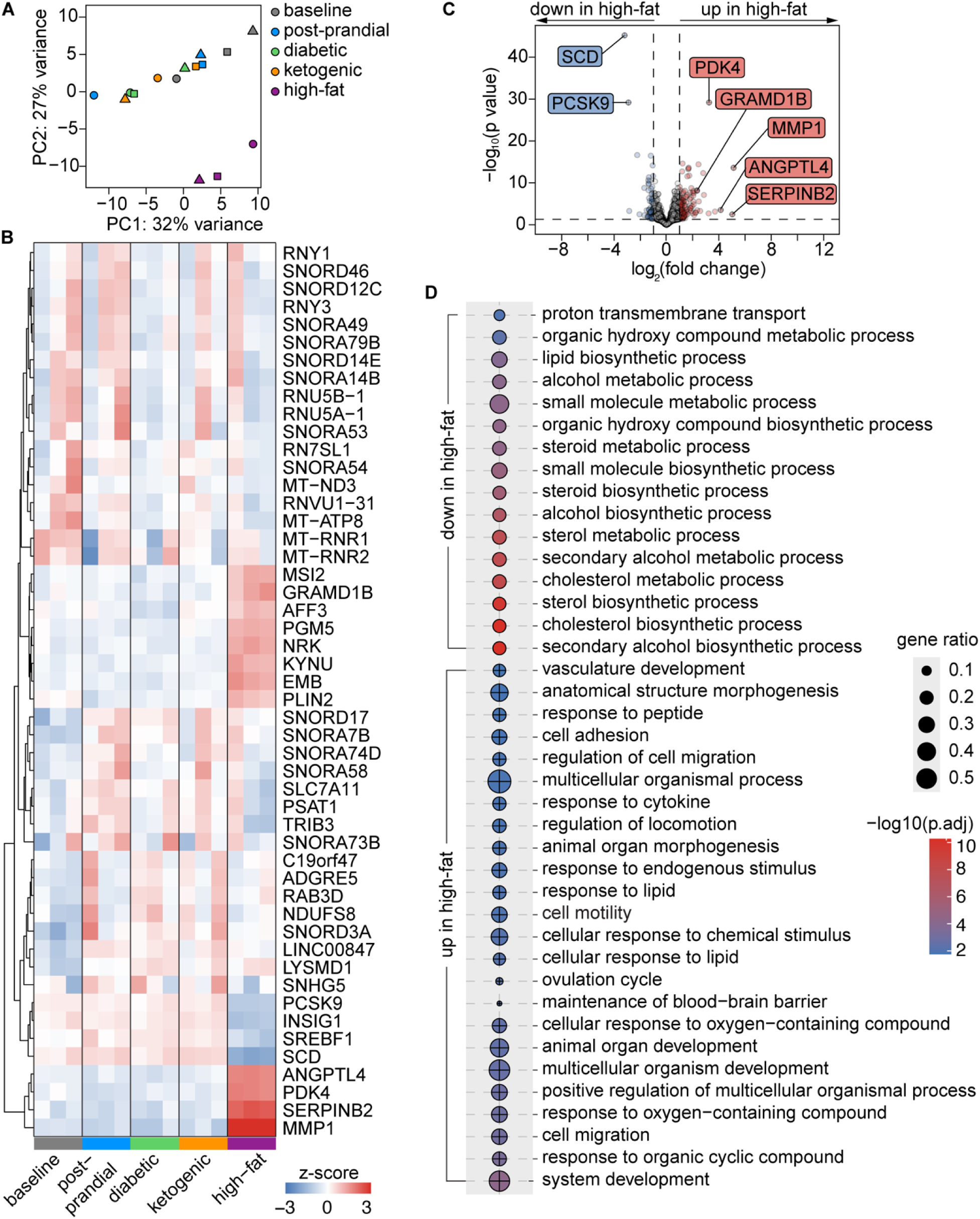
Transcriptomic analysis reveals gene-expression patterns specific to different circulating nutrient conditions. (**A**) Principal component analysis (PCA) of the 1000 most variable genes in cells cultured under five different media conditions. (**B**) Heatmap displaying hierarchical clustering of differentially expressed genes across the five media conditions. Each row represents a gene, and each column represents a condition. Red and blue indicate upregulated and downregulated genes, respectively, relative to the mean expression across conditions. (**C**) Volcano plot comparing gene expression between cells cultured under baseline and high-fat conditions. Genes with a fold change ≥ 2 or ≤-2 and adjusted p-value < 0.05 are highlighted in red (upregulated) and blue (downregulated), respectively. (**D**) Over-representation analysis with the 40 most significant GO terms of differentially expressed genes in cells cultured under high-fat compared to baseline condition. Each dot represents a biological process, with color representing statistical significance (-log10 adjusted p-value). Downregulated (circles) and upregulated (crossed circles) processes are displayed.

Consistently, gene-ontology analysis revealed significant downregulation of gene ontology (GO) terms related to lipid biosynthesis in cells cultured under high-fat conditions as compared to baseline (**Fig. 2D**). Additionally, GO terms associated with cellular migration and cell adhesion were upregulated by culture in high-fat media. These findings suggest that the circulating nutrients and metabolites characteristic of a high-fat diet might promote transcriptional changes that alter metabolism and invasive potential.

### High-fat conditions alter metabolic activity of MDA-MB-231 cells

To determine whether the different media conditions led to alterations in energy metabolism, we measured basal oxygen consumption rate (OCR) and extracellular acidification rate (ECAR) using a Seahorse extracellular flux analyzer (**Fig. S4A, B**). We found that the OCR of cells cultured under DMEM was ∼250 pmol/min/10^5^ cells (**Fig. S4C**), consistent with previous reports of MDA-MB-231 cells in their conventional (non-physiological) media^44,45^. Cells cultured under baseline, post-prandial, diabetic, and ketogenic conditions had an essentially identical OCR of ∼250 pmol/min/10^5^ cells (**Fig. 3A**), similar to that of cells cultured in DMEM. In contrast, cells cultured in high-fat media had an OCR of ∼350 pmol/min/10^5^ cells (**Fig. 3A**), which is ∼40% higher than those cultured in the other HPLM conditions, suggesting increased mitochondrial activity. However, we found no differences in the basal ECAR values of cells cultured in DMEM (**Fig. S4D**) or in the different HPLM formulations (**Fig. 3B**). Consistently, measurements of glucose concentration in conditioned media revealed that cells depleted glucose at similar rates (**Fig. S4E**). These observations suggest that high-fat conditions increase oxygen consumption but have no effect on glycolysis in MDA-MB-231 cells.

**Figure 3.**
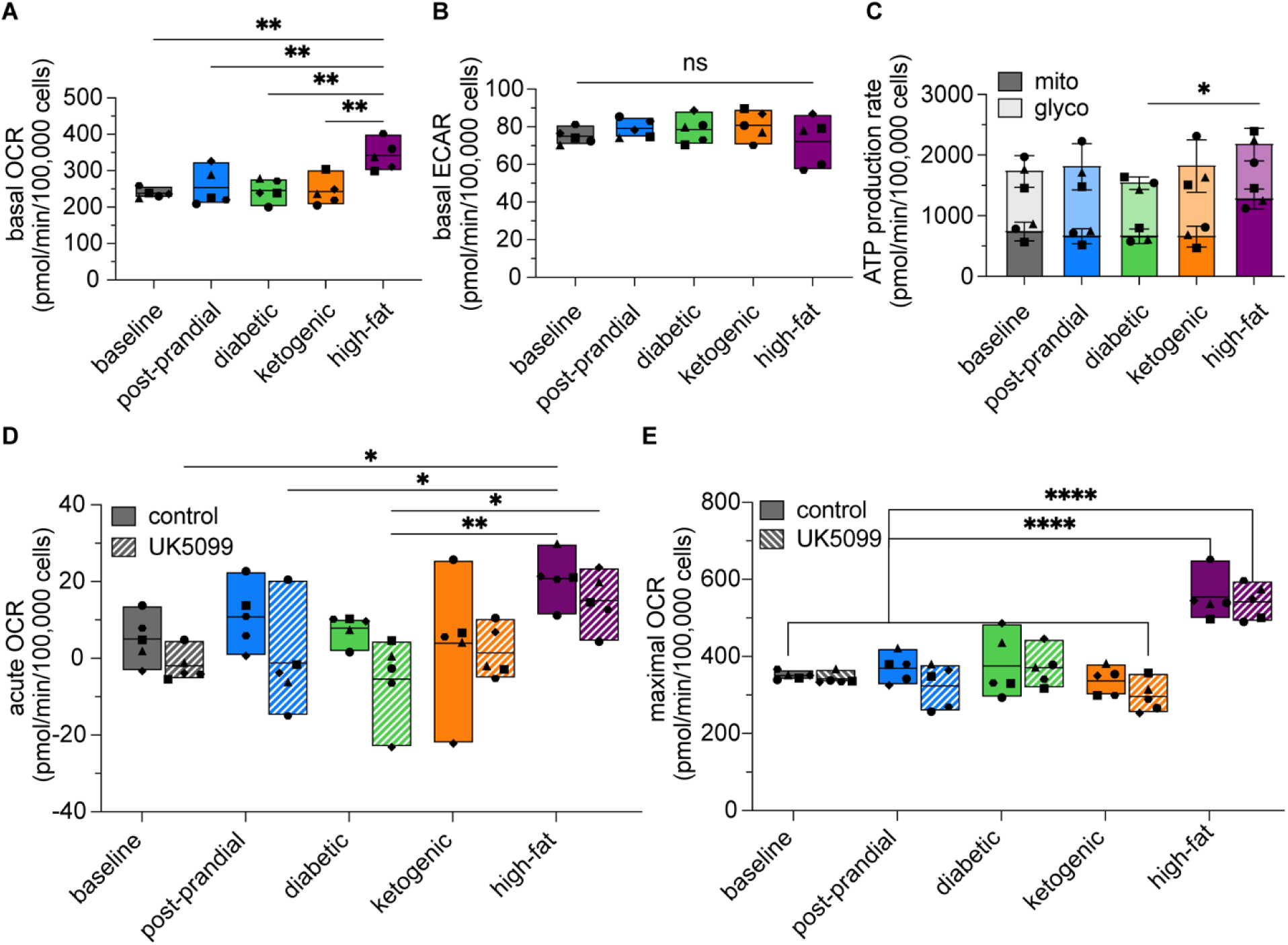
Cells cultured under high-fat conditions exhibit elevated metabolic activity. Graphs showing (**A**) basal oxygen consumption rate (OCR), (**B**) basal extracellular acidification rate (ECAR), and mitochondrial and glycolytic ATP-production rates (mitochondrial production rates: baseline 42.7%, post-prandial 36.6%, diabetic 43%, ketogenic 36%, and high-fat 58.6%; glycolytic production rates: baseline 57.3%, post-prandial 63.4%, diabetic 57%, ketogenic 63%, and high-fat 41.4%). Graphs showing acute response to UK5099 and (**E**) maximal respiration after being exposed to UK5099. Shown are mean and range, for five independent experiments, each with 3-4 statistical replicates. (*), p<0.05; (**), p<0.01; (****), p<0.0001; ns, not significant.

To test the validity of this hypothesis, we calculated the fraction of ATP produced through mitochondrial oxidative phosphorylation (OXPHOS) and glycolysis using the OCR and ECAR measurements. In normal cells, glucose is primarily metabolized through OXPHOS to produce ATP; however, cancer cells can shift their metabolism to favor glycolysis even in the presence of oxygen, a phenomenon known as the Warburg effect^46^. This metabolic reprogramming allows cancer cells to rapidly generate ATP and biosynthetic precursors needed for proliferation. Our measurements revealed that in MDA-MB-231 cells cultured in DMEM, ∼52% of the ATP is generated through glycolysis (**Fig. S4F**). When cultured in HPLM, glycolytic ATP production increased to 57% under baseline conditions, further rising to 64% under ketogenic conditions. In contrast, glycolytic ATP production decreased to ∼41% under high-fat conditions (**Fig. 3C**).

These findings indicate that MDA-MB-231 cells rely on glycolysis when cultured in HPLM but shift toward a greater dependence on OXPHOS under high-fat conditions.

We next assessed mitochondrial oxidation of glucose and pyruvate to further understand the metabolic reprogramming that might be induced by the different media conditions. To test whether the mitochondria are using glucose/pyruvate for respiration, we treated cells with UK5099, which inhibits the mitochondrial pyruvate carrier (MPC) to block the transport of pyruvate from the cytoplasm into the mitochondrial matrix, thus preventing its use in OXPHOS^47^. When cultured in DMEM, MDA-MB-231 cells responded to UK5099 by decreasing their OCR ∼33% (**Fig. S4G**), consistent with previous observations of these cells under similar conditions^44,48^. Surprisingly, however, MDA-MB-231 cells showed no response to UK5099 when cultured in HPLM (**Fig. 3D**). These data suggest that in physiologically relevant media conditions, mitochondrial function is independent of pyruvate transport via MPC. Nonetheless, the maximal respiration of cells cultured in high-fat media was higher than that of cells in the other media (**Fig. 3E, Fig. S4H**). High-fat conditions thus appear to enhance the capacity for OXPHOS, potentially due to an increased availability of fatty acids as a fuel source.

### High-fat conditions accelerate tumor growth and invasion

In contrast to the static microenvironment of 2D culture systems, cells *in vivo* are exposed to the flow of interstitial fluid, which circulates the nutrients that surround them. To replicate these interstitial fluid dynamics, we used an engineered microfluidic tumor model^28,30,31^ and investigated the effects of the different media conditions on tumor growth and cancer cell invasion (**Fig. 4A**). This 3D culture model consists of a solid, packed aggregate of cancer cells embedded within a type I collagen gel (**Fig. 4B**). After two days of culture, a hydrostatic pressure difference is applied to induce fluid flow at a rate similar to that of interstitial fluid observed *in vivo*^28,49^. Over time, the tumors grow, expanding in diameter and length, and cells invade into the surrounding collagen.

**Figure 4.**
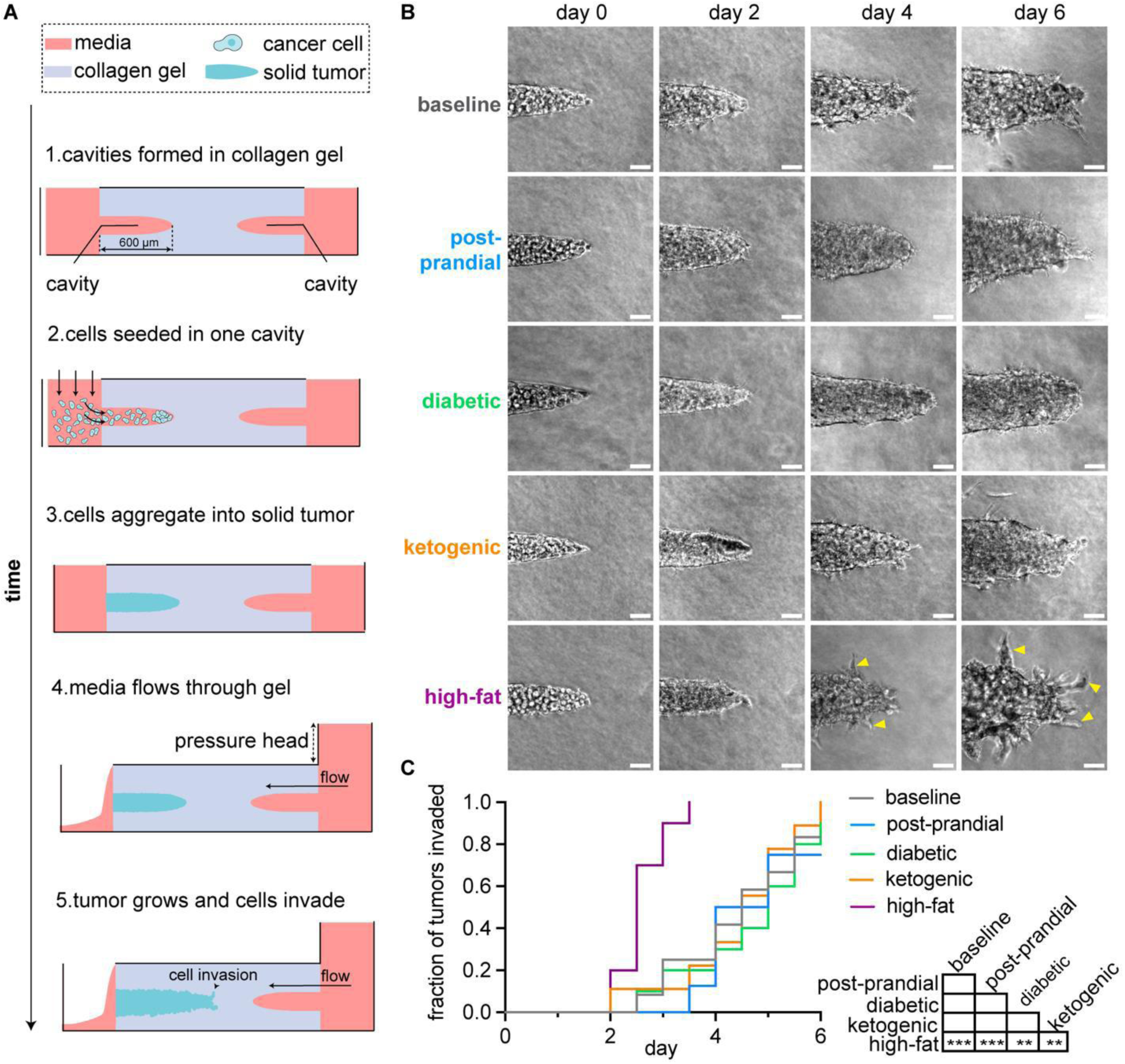
Tumors under high-fat conditions exhibit elevated rates of invasion. (**A**) Schematic of procedure for generating tumors cultured under interstitial fluid flow in microfluidic devices. (**B**) Phase-contrast images of tumors cultured under different media conditions taken on different days after seeding, showing 200 µm of the tumor tip. Arrow heads show point to example invasions; scale bars = 50 µm. (**C**) Kaplan-Meier plot of invasion from tumors cultured under different media conditions. Results are from 9-12 tumors in each condition, pooled across n = 3 independent experiments. (**), p<0.01; (***), p<0.001.

To investigate how circulating nutrients affect tumor metabolism, we performed mass spectrometry on conditioned media collected from engineered tumors cultured under the five different media conditions. We observed a consistent increase in the levels of several metabolites in each conditioned medium relative to its respective unconditioned control (**Fig. S5**).

Specifically, the conditioned media all contained increased levels of lactate and pyruvate. Similarly, we observed an increase in the levels of glutamate, suggesting enhanced glutaminolysis, a metabolic adaptation that helps sustain citric acid cycle activity by replenishing intermediates through increased glutamine metabolism^50^, and which is often observed in aggressive breast cancer cells. These findings are consistent with both the Warburg effect and our Seahorse data (**Fig. 3C**). The conditioned media were also depleted of hypoxanthine (**Fig. S5**), which may reflect an upregulation of the purine-salvage pathway to efficiently synthesize nucleotides^51^. Our analysis also revealed an increase in the levels of tryptophan, which promotes proliferation and invasion in MDA-MB-231 cells^52^ and may suggest limited catabolism through pathways such as the kynurenine pathway. Nonetheless, we surprisingly observed no differences in relative levels between the different media conditions.

To determine whether circulating nutrients influence functional behaviors such as tumor invasion, we monitored the tumors twice per day and noted when cells first invaded into the surrounding collagen. By day four, ∼40% of tumors in the baseline condition and ∼44% of the tumors in the post-prandial condition had invaded (**Fig. 4C**). Similarly, ∼30% of tumors in the diabetic condition and ∼33% of tumors in the ketogenic condition had invaded by this timepoint.

In contrast, the high-fat condition induced an accelerated rate of invasion, with 100% of these tumors invading by day 3.5. This observation is consistent with our gene-ontology analysis, which showed that culture in high-fat media leads to enrichment of GO terms related to cell adhesion and migration (**Fig. 2D**).

We also assessed whether circulating nutrients influence overall tumor growth by using a custom image-analysis pipeline to quantify different aspects of tumor morphology^27^ (**Fig. 5A**). We found that tumors cultured under high-fat conditions were significantly larger than those cultured under the other conditions (**Fig. 5B**). In contrast, tumors cultured under diabetic conditions were significantly smaller than those under baseline, ketogenic, or high-fat conditions (**Fig. 5B**). When we excluded the area occupied by invasions and segmented out the dense core of the tumors (**Fig. 5C**), we found that the tumor core metrics were roughly the same across all five conditions (**Fig. 5D-F**). However, the tumor cores under diabetic conditions were significantly smaller and narrower than those under both the post-prandial and high-fat conditions (**Fig. 5D, F**).

**Figure 5.**
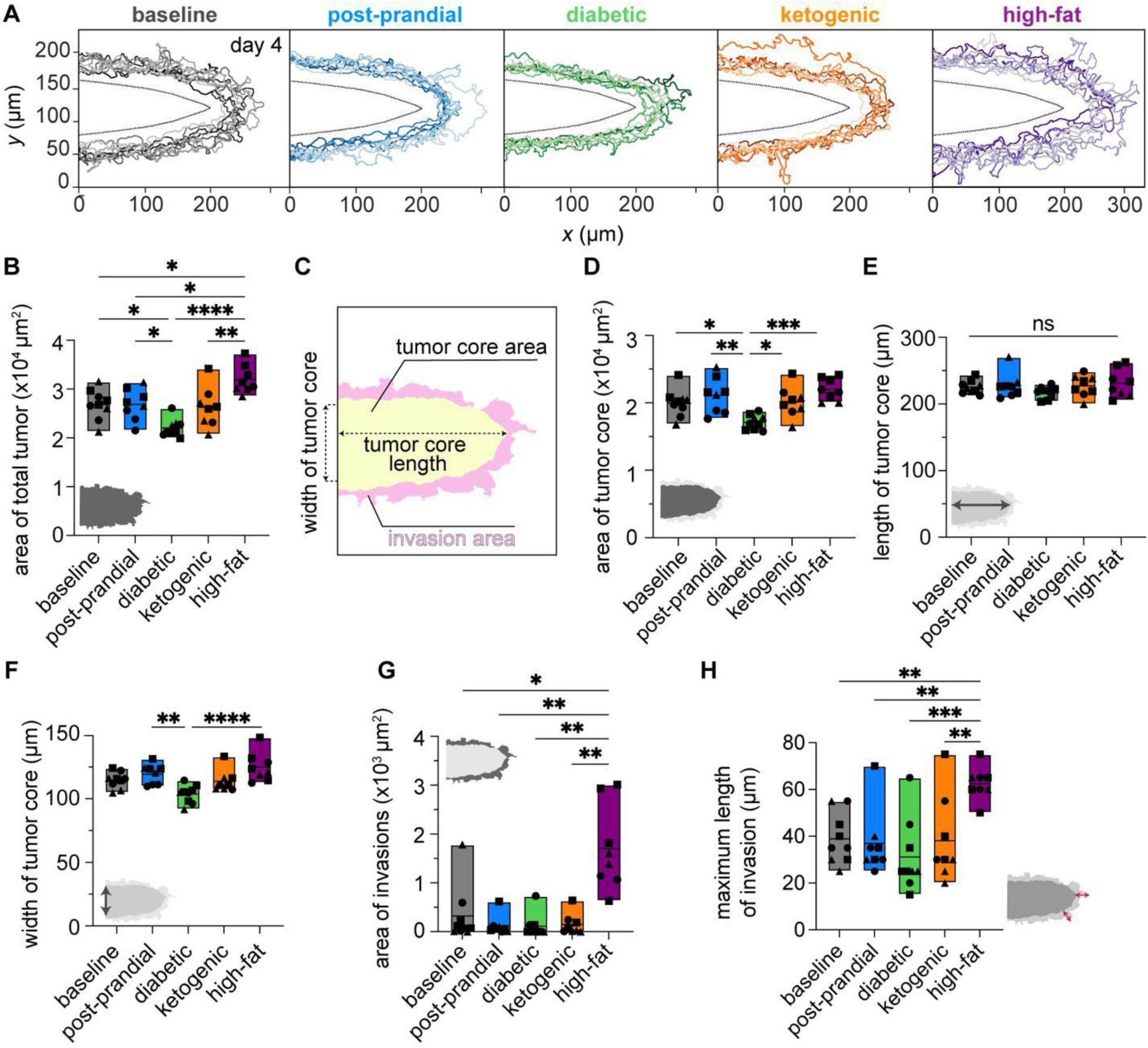
Tumors under high-fat conditions are larger in size. (**A**) Outlines of the segmented regions of tumors under each media condition, using images taken on day four. Each outline represents a different tumor within that condition. The outlines have been aligned at the cutoff point for segmentation and centered on this cutoff line on the left. (**B**) Graph of the area of the total tumor on day 4 after seeding across different media conditions. (**C**) Example of segmentation of the tumor core (yellow), width of the tumor core, and length of the tumor core. Graphs of the (**D**) area, (**E**) length, and (**F**) width of the tumor core across different media conditions. Graphs of the (**G**) area of invasions and (**H**) maximum length of invasion in different media conditions. Shown are mean and range for 8-9 tumors for each condition across n=3 independent experiments. (*), p<0.05; (**), p<0.01; (***), p<0.001; (****), p<0.0001; ns, not significant.

Additionally, we found that tumors under high-fat conditions had invasions that were significantly larger and longer than the other tumors (**Fig. 5G, H**), consistent with their accelerated rate of invasion (**Fig. 4B, C**). Overall, the high-fat conditions led to larger and more invasive tumors than the baseline, post-prandial, diabetic, or ketogenic conditions.

### High-fat conditions do not affect proliferation

Even though we did not detect a difference in proliferation across the media conditions in cells cultured in 2D, the elevated growth and invasion that we observed in tumors under high-fat conditions could still result from an increase in proliferation of the cancer cells in the 3D microenvironment under fluid flow. To test this possibility, we conducted EdU analysis on the engineered tumors (**Fig. 6A**). Despite the differences in tumor size and rates of invasion between tumors cultured under baseline and high-fat conditions, we found a similar number of cells (**Fig. 6B**) and rate of proliferation (**Fig. 6C**) across most conditions at the end of the experiment. The number of cells in tumors cultured under the diabetic condition was lower than other conditions, consistent with their smaller tumor area (**Fig. 5B**). Our RNA-sequencing analysis similarly revealed that the media conditions did not appreciably affect the expression of genes associated with proliferation (**Fig. S6A**). We therefore conclude that the increased size of tumors under high-fat conditions cannot be attributed to an increased rate of proliferation.

**Figure 6.**
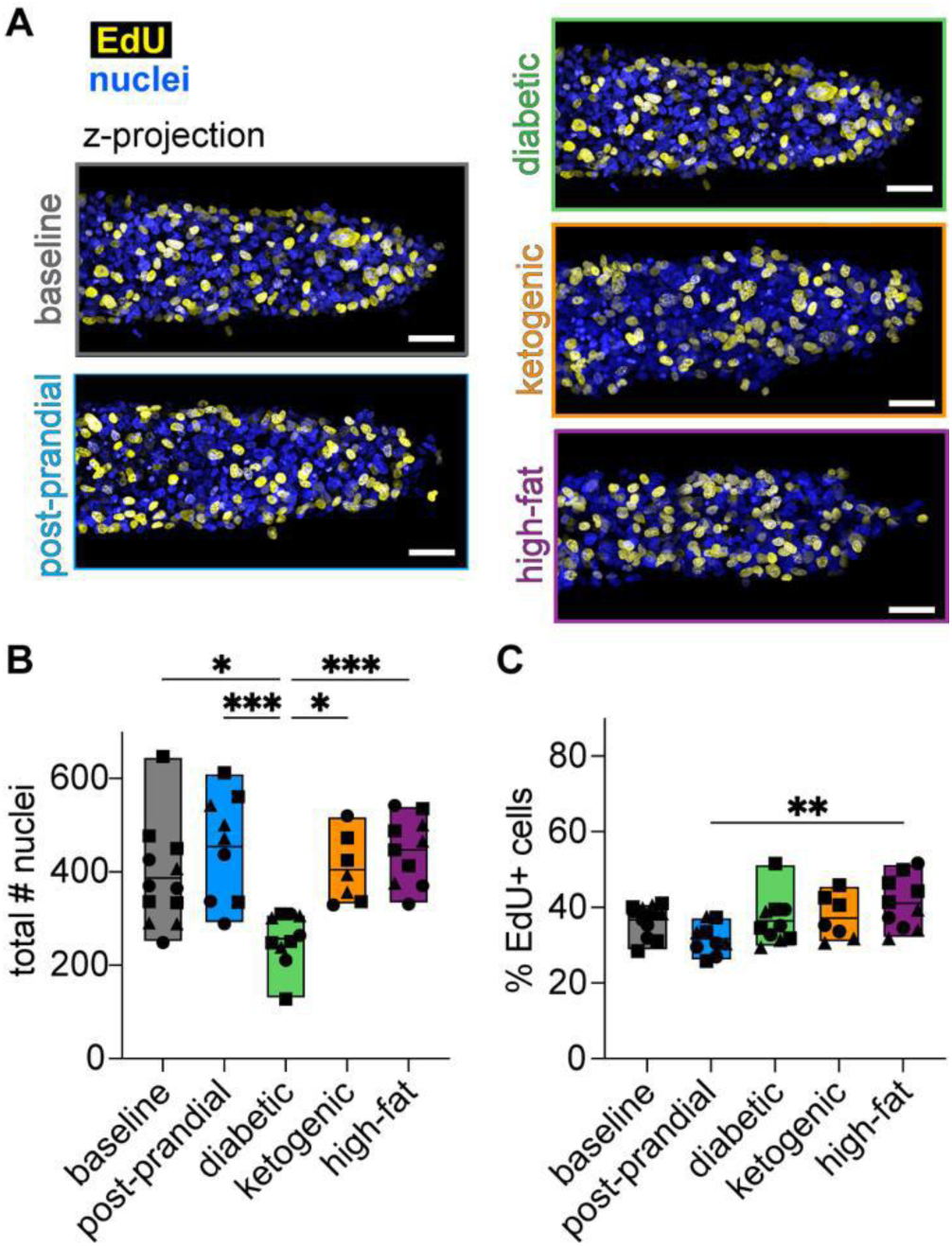
Tumors under high-fat conditions do not exhibit increased rates of proliferation. (**A**) Maximum-intensity z-projections of stacks of confocal images of EdU incorporation (yellow) in tumors under different media conditions, counterstained with Hoechst (blue) to visualize nuclei; scale bars = 50 µm. Graphs of the (**B**) total number of nuclei and (**C**) percentage of cells that incorporated EdU (EdU^+^ cells) in tumors under different media conditions. Shown are mean and range for 4-7 tumors in each media condition, pooled across n = 3 independent experiments. (*), p<0.05; (**), p<0.01; (***), p<0.001.

### High-fat conditions lead to formation of cavities within tumors

Given that the larger tumors observed under high-fat conditions are not linked to an increase in proliferation or number of cells, we investigated other potential causes. Upon examining z-stacks of nuclear stains, we observed that the centers of tumors under high-fat conditions lacked nuclei at the end of the experiment, while those in the baseline condition remained essentially solid (**Fig. 7A**). To confirm the presence of hollow regions in tumors under high-fat conditions and determine when they formed, we H&E stained tumors that were fixed and sectioned at different days of culture. This staining confirmed that hollow, cell-free cavities formed within tumors under high-fat conditions by day six, while the baseline controls retained a solid core (**Fig. 7B**).

**Figure 7.**
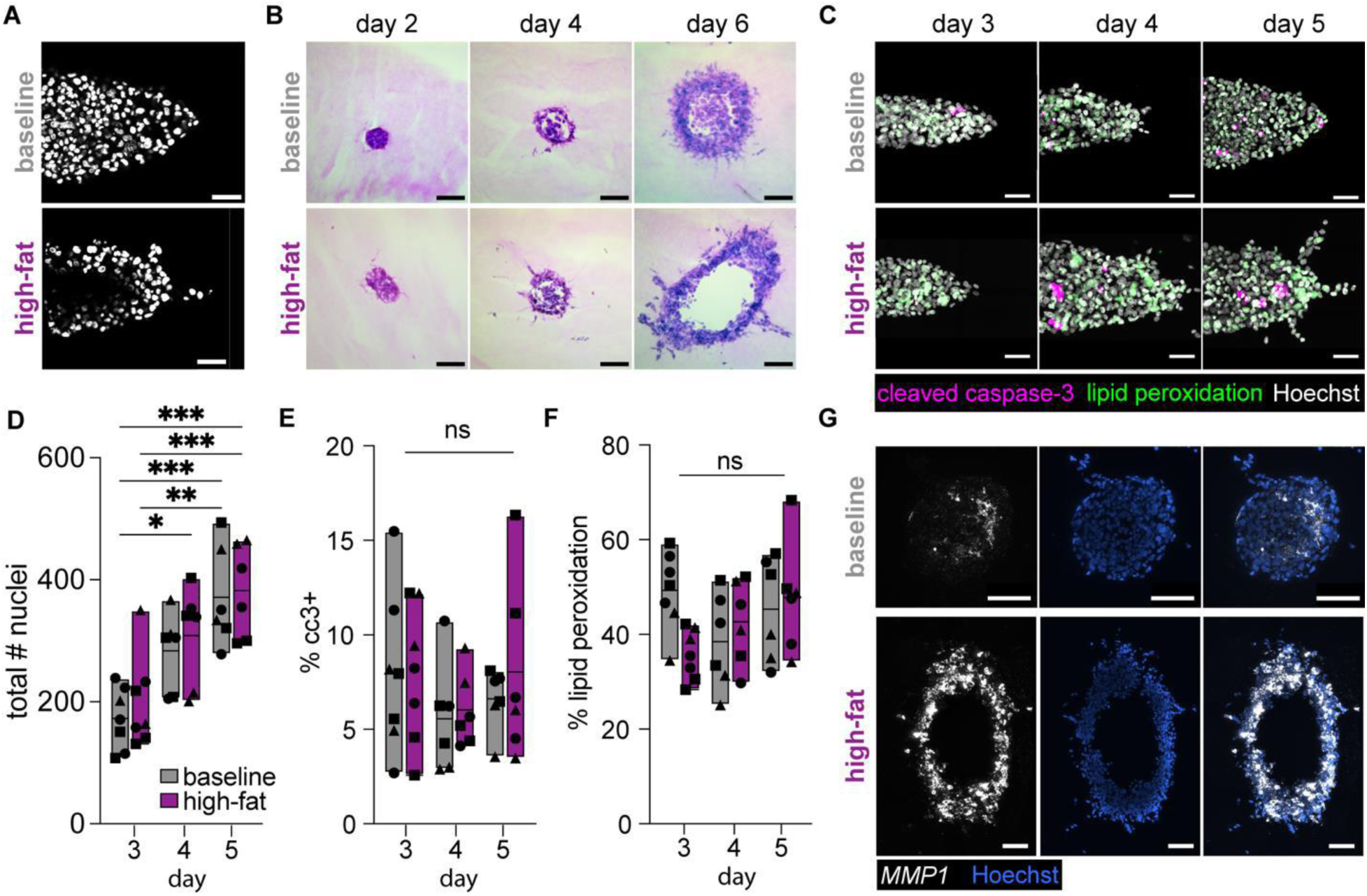
Tumors in high-fat conditions develop hollow regions that correlate with *MMP1* expression but not apoptosis or ferroptosis. (A) A z-slice from the center of DAPI-stained tumors in baseline (top) and high-fat (bottom) conditions on day six. (B) Images of H&E staining of tumor sections on days two, four, and six in baseline (top) and high-fat (bottom) conditions; scale bars = 100 𝜇m. (C) Representative images of tumors stained for cleaved caspase-3 (red), lipid peroxidation (green), and Hoechst (blue) on days three, four, and five in baseline (top) and high-fat (bottom) conditions; scale bars = 50 𝜇m. Graphs showing quantification of (**D**) total nuclei, (**E**) percentage of cleaved caspase-3^+^ cells (%cc3+), and (**F**) percentage of lipid peroxidation^+^ cells in tumors over time. Shown are mean and range for 6-7 tumors in each media condition, pooled across n = 3 independent experiments. (**G**) Expression of *MMP1* measured by RNAscope in tumor sections from baseline and high-fat conditions at day six of culture. (*), p<0.05; (**), p<0.01; (***), p<0.001; ns, not significant.

We hypothesized that the hollow regions might form because of cell death, as nutrient deprivation is known to induce apoptosis in the center of tumors^53^. To test this hypothesis, we fixed tumors at different time points and stained for both nuclei and cleaved-caspase-3 as a marker of apoptosis (**Fig. 7C**). We found that the total number of nuclei increased over time in both baseline and high-fat conditions (**Fig. 7D**), with no significant differences in cell number between the two conditions at any time point. Similarly, we observed no significant differences in the percentage of apoptotic cells between baseline and high-fat conditions (**Fig. 7E**).

Consistently, RNA-sequencing analysis revealed that cells under both conditions expressed low levels of apoptosis-related genes (**Fig. S6B**). These results suggest that hollowing of tumors under high-fat conditions occurs independently of apoptosis.

Although the circulating nutrient conditions did not result in differences in apoptosis, high-fat conditions might promote a different form of cell death. Specifically, an increase in the availability of fatty acids can enhance oxidative stress and promote the accumulation of peroxidized lipids within cells, which can lead to nonapoptotic cell death via ferroptosis^54^. To determine whether tumor hollowing resulted from ferroptosis, we measured lipid peroxidation in tumors cultured under baseline and high-fat conditions (**Fig. 7C**). Surprisingly, quantification of lipid peroxidation showed no significant differences between baseline and high-fat conditions (**Fig. 7F**). Since ferroptosis is an iron-dependent form of cell death, we also performed Prussian blue staining to assess iron accumulation within the tumors, which revealed neither detectable iron accumulation nor significant differences between baseline and high-fat conditions (**Fig. S7**). Consistent with the staining data, RNA-sequencing analysis showed that ferroptosis-related genes are expressed at low levels under both baseline and high-fat conditions (**Fig. S6C**). We therefore conclude that hollowing of high-fat tumors occurs independently of ferroptosis.

Our results show that high-fat tumors form hollow regions without increasing cell proliferation or death. The only other physical mechanism that could generate a hollow tumor is enhanced migration of cells away from the tumor core. This possibility is consistent with our RNA-sequencing data, which showed an increase in expression of genes associated with cell motility and ECM degradation, specifically in the high-fat condition (**Fig. 2B, C**). The gene most elevated in cells cultured in high-fat media is *MMP1*, an interstitial collagenase previously found to be upregulated in TNBC and associated with poor prognosis^55^. We therefore hypothesized that cells invading into the collagen surrounding high-fat tumors would show an increase in expression of *MMP1*. To test this hypothesis, we conducted fluorescence in situ hybridization (RNAscope) analysis. Our results confirmed that *MMP1* transcript is elevated in tumors cultured under high-fat conditions compared to baseline (**Fig. 7G**). Surprisingly, however, *MMP1* expression appears uniformly distributed throughout the tumor, with no clear spatial patterning associated with the invasive strands or core. These findings are consistent with the hypothesis that high-fat conditions enhance ECM degradation and invasion by leading to elevated expression of *MMP1*, thus leading to tumor hollowing.

## Discussion

Here, we adapted a microfluidic culture model to study the effects of circulating nutrients on tumor growth and cancer cell invasion. This reductionist approach allowed us to eliminate many of the complexities of *in vivo* conditions and focus solely on three key components of the TME: the ECM, the tumor, and the biochemical composition of the interstitial fluid. For the latter, we used human plasma-like medium^25^ to mimic five different circulating nutrient conditions: baseline, post-prandial, diabetic, ketogenic, and high-fat. RNA-sequencing analysis of cells cultured in HPLM under these five media conditions revealed a broad transcriptional reprogramming compared to those cultured in DMEM, highlighting the importance of using physiologically relevant media when studying the behavior of cancer cells in culture. Notably, cells cultured in the high-fat condition showed a transcriptomic shift, particularly in GO terms related to lipid metabolism and ECM reorganization. The upregulation of GO terms associated with lipid metabolism was expected, as the abundance of lipids in the high-fat media provides an ample energy source, prompting tumor cells to adapt their metabolism accordingly. Consistently, cells under high-fat conditions exhibited an increase in oxidative phosphorylation. The upregulation of GO terms associated with ECM reorganization was unexpected. These changes indicate an increased propensity for tissue remodeling and cell dissemination, which was not obvious in 2D culture. Consistently, however, tumors under high-fat conditions were larger, more invasive, and formed hollow regions over time, despite showing similar rates of proliferation, apoptosis, and ferroptosis as those under other conditions. Hollowing of these tumors correlated with elevated expression of the collagenase *MMP1* and increased cell invasion. Our results collectively highlight the impact of circulating nutrients on gene expression, metabolism, growth, and invasion, independent of alterations to the tumor genome.

These findings complement studies investigating the effects of high-fat diets in both cell-culture models and mouse models of cancer. In previous work, we found that adipose cells surrounding engineered tumors enhance the aggressiveness of MDA-MB-231 cells by secreting soluble factors that directly promote tumor cell invasion and vasculogenic escape^26,33^. In a mouse model comparing different diets, the largest tumors formed under a high-fat and moderate-carbohydrate diet^56^, which is analogous to the high-fat media used here. In addition, the elevated metabolic capacity that we observed in the high-fat condition might contribute to the observed increase in mitochondrial activity and could play a role in the aggressive behavior often associated with high-fat dietary states in cancer progression^57,58^.

Our results are consistent with emerging evidence that suggests that high-fat conditions can increase the expression of *MMP1* in cancer cells, including breast cancer. For example, MDA-MB-231 cells showed no changes in proliferation, but exhibited increased migration and elevated expression of *MMP1* when exposed to lipid-rich microenvironments^59^. A palmitate-rich microenvironment was recently found to promote metastasis in breast cancer by activating NF-κB signaling^60^, a pathway previously shown to regulate *MMP1* expression^61^. High concentrations of oleic acid were shown to promote *MMP1* expression and drive greater migration and invasion in endometrial cancer cells^62^. Although current evidence linking high-fat conditions to the upregulation of *MMP1* is limited, our findings provide direct support for this association in MDA-MB-231 cells, reinforcing the relevance of diet-induced transcriptional changes in breast cancer progression. Future studies will be needed to determine which fatty acids are inducing *MMP1* and whether similar transcriptional responses occur *in vivo*.

Several studies have suggested that elevated levels of glucose accelerate cancer cell proliferation^63^ and promote migration^64,65^, or that a ketogenic diet delays tumor growth^13,18,66^. However, we did not observe these effects in our diabetic and ketogenic conditions, either in 2D or 3D under flow. Instead, we found that tumors comprised of MDA-MB-231 cells cultured under diabetic conditions were smaller in overall size. It is important to note that our model is a simplified system that does not incorporate key components of the *in vivo* microenvironment, such as stromal cells, immune interactions, and systemic metabolic effects. These missing elements likely influence how cancer cells respond to circulating nutrients. For instance, the cytokine IL-22, which plays a key role in regulating immune responses and maintaining mucosal barrier integrity, was found to be reduced in both obese leptin receptor–deficient mice and in mice with hyperglycemia induced by a high-fat diet, compared to healthy control mice^67^.

Therefore, our findings suggest that, at least for the TNBC cell line used here, the post-prandial, diabetic, and ketogenic conditions may not directly influence the cancer cells themselves but might instead impact other features of the TME that are absent from our model. This possibility highlights the strength of using microfluidic models to disentangle cancer cell–intrinsic responses from those mediated by other components of the TME.

Previous studies have shown that MDA-MB-231 cells exhibit distinct gene-expression profiles depending on the composition of the culture medium^68^ and when they are exposed to fluid flow^69^. While DMEM remains a standard medium for cell culture, our RNA-sequencing analysis revealed that transitioning MDA-MB-231 cells into HPLM led to broad transcriptional reprogramming, with significant changes across several GO terms, including those related to metabolism, invasion, stress response, and tumor suppression. This observation suggests that standard culture media may obscure biologically relevant behaviors by failing to replicate the nutrient and ion composition of human plasma. Therefore, it is important to emphasize that combining a physiologically relevant biochemical microenvironment with interstitial fluid flow, both often overlooked in conventional models, is useful for more comprehensively assessing tumor cell behavior.

## Abbreviations

3D: three-dimensional
DEG: differentially expressed gene
ECAR: extracellular acidification rate
ECM: extracellular matrix
EMT: epithelial-to-mesenchymal transition
HPLM: human plasma-like medium
MMP1: matrix metalloproteinase 1
MPC: mitochondrial pyruvate carrier
OCR: oxygen consumption rate
OXPHOS: oxidative phosphorylation
TME: tumor microenvironment
TNBC: triple-negative breast cancer

## Acknowledgements

We thank Wei Wang and the Genomics Core Facility of Princeton University for assistance with transcriptomics, and Aaron Griffing for sharing his modified H&E staining protocol. This work was supported in part by grants from the National Institutes of Health (CA187692, CA214292) and the Ludwig Institute for Cancer Research Princeton. M.K. was supported in part by the National Science Foundation (through the Center for the Physics of Biological Function PHY-1734030) and the National Cancer Institute (through Rutgers Cancer Institute of New Jersey 1T32CA257957). C.T.-Y. was supported in part by the Damon Runyon Cancer Research Foundation through the 2023 Damon Runyon Quantitative Biology Fellowship (DRQ-17-23). M.C.B-S. was supported in part by a New Jersey Commission on Cancer Research Predoctoral Fellowship (COCR22PRF009).

## Notes

### Competing Interest Statement

The authors have declared no competing interest.

## References

1 Fujisaka, S. et al. Diet, genetics, and the gut microbiome drive dynamic changes in plasma metabolites. Cell Rep. 22, 3072–3086 (2018).

2 Pellis, L. et al. Plasma metabolomics and proteomics profiling after a postprandial challenge reveal subtle diet effects on human metabolic status. Metabolomics 8, 347–359 (2012).

3 Qian, L., Zhang, F., Yin, M. & Lei, Q. Cancer metabolism and dietary interventions. Cancer Biol. Med. 19, 163–174 (2021).

4 Mittelman, S. D. The role of diet in cancer prevention and chemotherapy efficacy. Annu. Rev. Nutr. 40, 273–297 (2020).

5 O’Keefe, S. J. D. Diet, microorganisms and their metabolites, and colon cancer. Nat. Rev. Gastroenterol. Hepatol. 13, 691–706 (2016).

6 Taylor, S. R., Falcone, J. N., Cantley, L. C. & Goncalves, M. D. Developing dietary interventions as therapy for cancer. Nat. Rev. Cancer 22, 452–466 (2022).

7 Rossi, R. E., Pericleous, M., Mandair, D., Whyand, T. & Caplin, M. E. The role of dietary factors in prevention and progression of breast cancer. Anticancer Res. 34, 6861–6876 (2014).

8 Lien, E. C. & Vander Heiden, M. G. A framework for examining how diet impacts tumour metabolism. Nat. Rev. Cancer 19, 651–661 (2019).

9 Tang, L., Qu, R. W., Park, J., Simental, A. A. & Inman, J. C. Prevalence of occult central lymph node metastasis by tumor size in papillary thyroid carcinoma: a systematic review and meta-analysis. Curr. Oncol. 30, 7335–7350 (2023).

10 Emond, J. A. et al. Risk of breast cancer recurrence associated with carbohydrate intake and tissue expression of IGFI receptor. CEBP 23, 1273–1279 (2014).

11 Ho, V. W. et al. A low carbohydrate, high protein diet slows tumor growth and prevents cancer initiation. Cancer Res. 71, 4484–4493 (2011).

12 Yakar, S. et al. Increased tumor growth in mice with diet-induced obesity: impact of ovarian hormones. Endocr. J. 147, 5826–5834 (2006).

13 Yang, L. et al. Ketogenic diet and chemotherapy combine to disrupt pancreatic cancer metabolism and growth. Med 3, 119–136 (2022).

14 Ringel, A. E. et al. Obesity shapes metabolism in the tumor microenvironment to suppress anti-tumor immunity. Cell 183, 1848–1866 e1826 (2020).

15 Tang, F. Y., Pai, M. H. & Chiang, E. P. Consumption of high-fat diet induces tumor progression and epithelial-mesenchymal transition of colorectal cancer in a mouse xenograft model. J. Nutr. Biochem 23, 1302–1313 (2012).

16 Sundram, S. Y., Lin;. High-fat diet enhances mammary tumorigenesis and pulmonary metastasis and alters inflammatory and angiogenic profiles in MMTV-PyMT mice. Anticancer Res. 36, 6279–6288 (2016).

17 Dogan, S., et al. Effects of high-fat diet and/or body weight on mammary tumor leptin and apoptosis signaling pathways in MMTV-TGF-α mice. BCR 9, R91 (2007).

18 Grube, M. et al. Ketogenic diet does not promote triple-negative and luminal mammary tumor growth and metastasis in experimental mice. Clin. Exp. Metastasis 41, 251–266 (2024).

19 Su, Z. L., Yanqing; Xia, Zhangchuan; Rustgi, Anil K.; Gu, Wei;. An unexpected role for the ketogenic diet in triggering tumor metastasis by modulating BACH1-mediated transcription. Sci. Adv. 10 (2024).

20 Fine, E. Z., Yiyu; Koba, Wade;. FDG-PET of mouse breast cancers on ketogenic vs. standard chow diets, with or without added rapamycin. J. Nucl. Med. 60 (2019).

21 Nathanson, S. D. & Nelson, L. Interstitial fluid pressure in breast cancer, benign breast conditions, and breast parenchyma. Ann. Surg. Oncol. 1, 333–338 (1994).

22 Butler, T. P., Grantham, F. H. & Gullino, P. M. Bulk transfer of fluid in the interstitial compartment of mammary tumors. Cancer Res. 35, 3084–3088 (1975).

23 Brennan, M. C. & Nelson, C. M. in Biomaterial Based Approaches to Study the Tumour Microenvironment (eds J. O. Winter & S Rao) Ch. 7, 163–196 (Royal Society of Chemistry, 2022).

24 Benien, P. & Swami, A. 3D tumor models: history, advances and future perspectives. Future Oncol. 10, 1311–1327 (2014).

25 Cantor, J. R. et al. Physiologic medium rewires cellular metabolism and reveals uric acid as an endogenous inhibitor of UMP synthase. Cell 169, 258–272 e217 (2017).

26 Dance, Y. W. et al. Adipose cells induce escape from an engineered human breast microtumor independently of their obesity status. Cell. Mol. Bioeng. 16, 23–39 (2023).

27 Leggett, S. E., Brennan, M. C., Martinez, S., Tien, J. & Nelson, C. M. Relatively rare populations of invasive cells drive progression of heterogeneous tumors. Cell. Mol. Bioeng. (2024).

28 Tien, J., Dance, Y. W., Ghani, U., Seibel, A. J. & Nelson, C. M. Interstitial hypertension suppresses escape of human breast tumor cells via convection of interstitial fluid. Cell. Mol. Bioeng. 14, 147–159 (2021).

29 Rabie, E. M. et al. Matrix degradation and cell proliferation are coupled to promote invasion and escape from an engineered human breast microtumor. Integr. Biol. 13, 17–29 (2021).

30 Piotrowski-Daspit, A. S., Tien, J. & Nelson, C. M. Interstitial fluid pressure regulates collective invasion in engineered human breast tumors via Snail, vimentin, and E-cadherin. Integr. Biol. 8, 319–331 (2016).

31 Piotrowski-Daspit, A. S., Simi, A. K., Pang, M., Tien, J. & Nelson, C. M. in Mammary Gland Development: Methods and Protocols Vol. 1501 *Methods in Molecular Biology* (eds F Martin, T Stein, & J Howlin) 245–257 (SpringerLink, 2017).

32 Seibel, A. J., Kelly, O. M., Dance, Y. W., Nelson, C. M. & Tien, J. Role of lymphatic endothelium in vascular escape of engineered human breast microtumors. Cell. Mol. Bioeng. 15, 553–569 (2022).

33 Dance, Y. W. et al. Adipose stroma accelerates the invasion and escape of human breast cancer cells from an engineered microtumor. Cell. Mol. Bioeng. 15, 15–29 (2022).

34 Love, M. I., Huber, W. & Anders, S. Moderated estimation of fold change and dispersion for RNA-seq data with DESeq2. Genome Biol. 15 (2014).

35 Zhu, A., Ibrahim, J. G. & Love, M. I. Heavy-tailed prior distributions for sequence count data: removing the noise and preserving large differences. Bioinform. 35, 2084–2092 (2019).

36 Kolberg, L., Raudvere, U., Kuzmin, I., Vilo, J. & Peterson, H. gprofiler2--an R package for gene list functional enrichment analysis and namespace conversion toolset g:Profiler. F1000Res. 9, 709 (2020).

37 Humason, G. Animal Tissue Techniques. 661 (WH Freeman and Company, 1979).

38 Hanahan, D. Hallmarks of cancer: new dimensions. Cancer Discov. 12, 31–46 (2022).

39 Bera, K. et al. Extracellular fluid viscosity enhances cell migration and cancer dissemination. Nature 611, 365–373 (2022).

40 Naghedi-Baghdar, H. et al. Effect of diet on blood viscosity in healthy humans: a systematic review. Electron. Physician 10, 6563–6570 (2018).

41 Liu, C.-Y., Lin, H.-H., Tang, M.-J. & Wang, Y.-K. Vimentin contributes to epithelial-mesenchymal transition cancer cell mechanics by mediating cytoskeletal organization and focal adhesion maturation. Oncotarget 6, 15966–15983 (2015).

42 Chénais, B., Cornec, M., Dumont, S., Marchand, J. & Blanckaert, V. Transcriptomic response of breast cancer cells MDA-MB-231 to docosahexaenoic acid: downregulation of lipid and cholesterol metabolism genes and upregulation of genes of the pro-apoptotic ER-stress pathway. Int. J. Environ. Res. Public Health 17, 3746 (2020).

43 Kersten, S. Role and mechanism of the action of angiopoietin-like protein ANGPTL4 in plasma lipid metabolism. J. Lipid Res. 62, 100150 (2021).

44 Winter, M. et al. Mitochondrial adaptation decreases drug sensitivity of persistent triple negative breast cancer cells surviving combinatory and sequential chemotherapy. Neoplasia 46, 100949 (2023).

45 Phan, L. et al. The cell cycle regulator 14-3-3σ opposes and reverses cancer metabolic reprogramming. Nat. Commun. 6, 7530 (2015).

46 Warburg, O., Wind, F. & Negelein, E. The metabolism of tumors in the body. J Gen Physiol. 8.6, 519 (1927).

47 Christen, S. et al. Breast cancer-derived lung metastases show increased pyruvate carboxylase-dependent anaplerosis. Cell Rep. 17, 837–848 (2016).

48 Carreira, A. S. A. et al. Mitochondrial rewiring drives metabolic adaptation to NAD(H) shortage in triple negative breast cancer cells. Neoplasia 41 (2023).

49 Chary, S. R. & Jain, R. K. Direct measurement of interstitial convection and diffusion of albumin in normal and neoplastic tissues by fluorescence photobleaching. PNAS 86, 5385–5389 (1989).

50 Erickson, J. W. & Cerione, R. A. Glutaminase: A hot spot for regulation of cancer cell metabolism? Oncotarget 1, 734–740 (2010).

51 Tran, D. H. et al. De novo and salvage purine synthesis pathways across tissues and tumors. Cell 187, 3602–3618.e3620 (2024).

52 Gautam, J. et al. Tryptophan hydroxylase 1 and 5-HT7 receptor preferentially expressed in triple-negative breast cancer promote cancer progression through autocrine serotonin signaling. Mol. Cancer 15 (2016).

53 Chen, Z., Han, F., Du, Y., Shi, H. & Zhou, W. Hypoxic microenvironment in cancer: molecular mechanisms and therapeutic interventions. Sig. Transduct. Target Ther. 8 (2023).

54 Dai, E. et al. A guideline on the molecular ecosystem regulating ferroptosis. Nat. Cell Biol. 26, 1447–1457 (2024).

55 Westermarck, J. & Kähäri, V. M. Regulation of matrix metalloproteinase expression in tumor invasion. ASEB J. 13, 781–792 (1999).

56 Mavropoulos, J. C. et al. The effects of varying dietary carbohydrate and fat content on survival in a murine LNCaP prostate cancer xenograft model. CAPR 2, 557–565 (2009).

57 Zhao, Y., et al. Pubertal high fat diet: effects on mammary cancer development. BCR 15, R100 (2013).

58 Kim, E. J., et al. Dietary fat increases solid tumor growth and metastasis of 4T1 murine mammary carcinoma cells and mortality in obesity-resistant BALB/c mice. BCR 13, R78 (2011).

59 Koellensperger, E. et al. The impact of human adipose tissue-derived stem cells on breast cancer cells: implications for cell-assisted lipotransfers in breast reconstruction. Stem Cell Res. Ther. 8 (2017).

60 Altea-Manzano, P. et al. A palmitate-rich metastatic niche enables metastasis growth via p65 acetylation resulting in pro-metastatic NF-κB signaling. *Nat*. Cancer 4, 344–364 (2023).

61 Nguyen, C. H. et al. NF-κB contributes to MMP1 expression in breast cancer spheroids causing paracrine PAR1 activation and disintegrations in the lymph endothelial barrier in vitro. Oncotarget 6, 39262–39275 (2015).

62 Feng, J. et al. Oleic acid promotes the development of endometrial cancer by up-regulating KLF4 expression. Eur. J. Gynaecol. Oncol. 41, 432–438 (2020).

63 Bao, Z. et al. High glucose promotes human glioblastoma cell growth by increasing the expression and function of chemoattractant and growth factor receptors. Transl. Oncol. 12, 1155–1163 (2019).

64 Phoomak, C. et al. High glucose levels boost the aggressiveness of highly metastatic cholangiocarcinoma cells via O-GlcNAcylation. Sci. Rep. 7, 43842 (2017).

65 Azoulay, S. et al. KIT is highly expressed in adenoid cystic carcinoma of the breast, a basal-like carcinoma associated with a favorable outcome. Mod. Pathol. 18, 1623–1631 (2005).

66 Otto, C. et al. Growth of human gastric cancer cells in nude mice is delayed by a ketogenic diet supplemented with omega-3 fatty acids and medium-chain triglycerides. BMC Cancer 8, 122 (2008).

67 Wang, X. et al. Interleukin-22 alleviates metabolic disorders and restores mucosal immunity in diabetes. Nature 514, 237–241 (2014).

68 Kim, S. W., Kim, S.-J., Langley, R. R. & Fidler, I. J. Modulation of the cancer cell transcriptome by culture media formulations and cell density. Int. J. Oncol. 46, 2067–2075 (2015).

69 Fuh, K. F., Shepherd, R. D., Withell, J. S., Kooistra, B. K. & Rinker, K. D. Fluid flow exposure promotes epithelial-to-mesenchymal transition and adhesion of breast cancer cells to endothelial cells. BCR 23 (2021).

