## Supplemental material for "High-fat circulating nutrients promote growth and invasion in a 3D microfluidic tumor model of triple-negative breast cancer"

### Supplementary Information

### Supplementary Text

To uncover possible changes in gene expression when cells are transitioned from DMEM into baseline HPLM, we performed RNA-sequencing analysis of MDA-MB-231 cells cultured in 2D (**Fig. S2**). We found that culture in HPLM increases the expression of *EPHB6*, a kinase-deficient Eph receptor known to suppress metastasis in breast cancer[1] that is often silenced in aggressive breast cancer cells in standard culture[2]. In addition, culture in HPLM increases the expression of *KRT8*, an epithelial cytokeratin associated with a decrease in proliferation, motility, and metastasis in breast cancer cell lines[3]. HPLM also increases the expression of *TNFSF15*, a member of the tumor necrosis factor (TNF) superfamily that is linked to endothelial cell apoptosis and inhibits tumor angiogenesis[4]. Consistently, HPLM enhances the expression of *MYCT1*, a Myc target that has been shown to impede cancer cell migration[5]. The induction of these genes, in addition to *ZBTB7C*, which has been shown to have tumor-suppressor effects[6], suggests that cells cultured in HPLM might adopt a less aggressive phenotype than those cultured in DMEM. On the other hand, we also found that culture in HPLM decreases the expression of *CA2*, *B3GNT7*, and *BEND4*, suggesting a more complex picture. Specifically, *CA2*, which encodes carbonic anhydrase II, is a key enzyme involved in coordinating pH-sensitive signaling pathways that promote cancer cell survival and proliferation[7,8]. In addition, *B3GNT7* is a glycosyltransferase implicated in modulating glycosylation patterns that influence cell proliferation, migration, and invasion; its loss has been linked to decreased proliferation, migration, and invasion in breast cancer models[9]. Consistently, *BEND4* encodes BEN domain-containing protein 4 and plays a role in tumor suppression and cellular stress responses[10].

Finally, GO analysis reveals that culture in HPLM leads to significant upregulation of GO terms involved in immune response, cytokine signaling, and ECM organization (**Fig. S3A**). Altogether, our data suggest substantial shifts in gene expression when cells are cultured in HPLM.

To investigate the transcriptional impact of our five different dietary conditions, we performed a PCA analysis. PCA of the 1000 most variable genes revealed that cells cultured in post-prandial, diabetic, and ketogenic media clustered with those in baseline medium (**Fig. 2A** and **Fig. S2A**), suggesting overall similarity in global gene-expression profiles. Nevertheless, pairwise comparisons of each condition with the baseline identified subsets of differentially expressed genes, indicating condition-specific transcriptional responses (**Fig. 2B**). Here, we highlight the most differentially expressed genes under each of the post-prandial, diabetic, and ketogenic conditions, focusing on those implicated in cancer-related processes.

A comparison between the baseline and post-prandial conditions reveals several differentially expressed genes (**Fig. S3B**). Specifically, *LSM3* is downregulated in the post-prandial condition. *LSM3* encodes a core component of the RNA-processing machinery involved in mRNA splicing and degradation. Its downregulation in this condition suggests a shift in RNA metabolism, where cells may reduce reliance on stress-induced mRNA storage and decay mechanisms in favor of active protein synthesis[11]. In addition, *APBA2*, a gene involved in cellular signaling and vesicle trafficking that has been implicated in regulating cancer cell motility, is downregulated in the post-prandial condition[12]. On the other hand, post-prandial conditions were found to upregulate *R3HDM4*, which encodes a nucleic acid-binding protein localized to the nucleus. Similarly, *SRGAP2B*, which encodes a Rho GTPase-activating protein paralog that may regulate cell migration and differentiation by inhibiting the full-length SRGAP2, was upregulated[13]. Related SRGAP-family proteins have been implicated in promoting cancer cell invasion and metastasis[14,15]. *CYTOR*, a long non-coding RNA known to promote EMT and metastasis across various cancer types, also showed increased expression under the post-prandial condition[16]. Finally, *COX6CP14*, a pseudogene of the mitochondrial cytochrome c oxidase subunit 6C, does not encode a protein but its increased expression in the post-prandial condition suggests a regulatory role, since pseudogene transcripts can act as microRNA sponges and are often dysregulated in tumors[17,18]. Taken together, these transcriptional changes indicate that the post-prandial biochemical microenvironment may promote gene-expression profiles associated with enhanced cellular adaptability, increased metabolic activity, and greater metastatic potential in breast cancer cells.

Comparison between the baseline and diabetic condition revealed that the latter upregulated *METTL4*, *R3HDM4*, *MIR545*, and *LINC01775*, while downregulating *MDK*, *KISS1*, *LSM3*, and *SLC25A5* (**Fig. S3C**). *METTL4* is an N6-adenine methyltransferase that adds epigenetic RNA/DNA modifications, and its increased expression can drive tumor metastasis under hypoxia[19]. The microRNA *MIR545* regulates cell cycle and DNA repair and has been implicated as a tumor-related miRNA in breast cancer[20]. *LINC01775* is a long noncoding RNA observed to be elevated in malignancies[21]. *MDK* encodes a heparin-binding growth factor that normally drives oncogenic traits like proliferation, migration, and angiogenesis, and is typically overexpressed in various cancers[22]. *KISS1* is a well-characterized metastasis-suppressor gene in breast cancer, and its reduced expression in diabetic conditions may indicate a microenvironment that favors tumor progression[23]. Additionally, downregulation of *LSM3* suggests potential shifts in metabolic gene regulation[11]. *SLC25A5* encodes the mitochondrial ADP/ATP carrier ANT2, which supports high glycolytic metabolism and its silencing triggers apoptosis in breast cancer[24]. Collectively, these transcriptional alterations suggest that diabetic conditions may reprogram breast cancer cells toward enhanced adaptability, potentially favoring tumor progression.

The ketogenic condition exhibited fewer transcriptional changes as compared to baseline, with no significantly downregulated genes, but an upregulation of *R3HDM4*, *SRGAP2B*, *LINC00847*, *MTMR11*, and *SNORD3A* (**Fig. S3D**). *R3HDM4* encodes a nuclear protein that enables nucleic acid binding activity, and altered expression of this gene has been observed in pan-cancer analyses[25]. Upregulation of *SRGAP2B* indicates a possible effect on metastatic behavior under ketogenic conditions[15]. *LINC00847* is transcribed as a long RNA without protein-coding capacity, and it is overexpressed in multiple cancers, including breast cancer, with high levels correlating to poorer patient prognosis[26,27]. *MTMR11* encodes a myotubularin-related lipid phosphatase predicted to act on phosphatidylinositol-3-phosphate and regulate phosphoinositide signaling. HER2-driven upregulation of *MTMR11* has been shown to promote tumor cell proliferation[28]. *SNORD3A* encodes the U3 small nucleolar RNA, a key component of the ribosome biogenesis machinery that guides pre-rRNA processing[29]. Aberrant expression of this non-coding RNA has been observed in breast cancer[30]. Taken together, these gene-expression changes indicate that the ketogenic condition may enhance cellular behaviors linked to metastasis and proliferation by modulating RNA biogenesis and lipid metabolism, highlighting distinct metabolic and regulatory adjustments.

**Supplementary References**


1 Truitt, L., Freywald, T., Decoteau, J., Sharfe, N. & Freywald, A. The EphB6 receptor cooperates with c-Cbl to regulate the behavior of breast cancer cells. *Cancer Res.* **70**, 1141-1153 (2010).

2 Bhushan, L. & Kandpal, R. P. EphB6 receptor modulates micro RNA profile of breast carcinoma cells. *PloS one* **6**, e22484 (2011).

3 Iyer, S. V. *et al.* Understanding the role of keratins 8 and 18 in neoplastic potential of breast cancer derived cell lines. *PloS one* **8**, e53532 (2013).

4 Zhang, Z. & Li, L.-Y. TNFSF15 modulates neovascularization and inflammation. *Cancer Microenviron.* **5**, 237-247 (2012).

5 Xu, J., Sun, Y., Fu, W. & Fu, S. MYCT1 in cancer development: Gene structure, regulation, and biological implications for diagnosis and treatment. *Biomed. Pharmacother.* **165** (2023).

6 Chen, X., Jiang, Z., Wang, Z. & Jiang, Z. The prognostic and immunological effects of ZBTB7C across cancers: friend or foe? *Aging* **13**, 12849-12864 (2021).

7 Raum, H. N., Fisher, S. Z. & Weininger, U. Energetics and dynamics of the proton shuttle of carbonic anhydrase II. *CMLS* **80** (2023).

8 Hulikova, A., Aveyard, N., Harris, A. L., Vaughan-Jones, R. D. & Swietach, P. Intracellular carbonic anhydrase activity sensitizes cancer cell pH signaling to dynamic changes in CO2 partial pressure. *J. Biol. Cell* **289**, 25418-25430 (2014).

9 Wang, X. *et al.* Identification of glycogene-based prognostic signature and validation of B3GNT7 as a potential biomarker and therapeutic target in breast cancer. *J. Cancer Res. Clin. Oncol.* **149**, 16957-16969 (2023).

10 Yao, Y. *et al.* Epigenetic silencing of BEND4, a novel DNA damage repair gene, is a synthetic lethal marker for ATM inhibitor in pancreatic cancer. *Front. Med.* **18**, 721-734 (2024).

11 Ta, H. D. K. *et al.* Potential therapeutic and prognostic values of LSM family genes in breast cancer. *Cancers* **13**, 4902 (2021).

12 Stoletov, K., Willetts, L., Beatty, P. H. & Lewis, J. D. Intravital imaging tumor screen used to identify novel metastasis-blocking therapeutic targets. *CES* **2**, 275-278 (2018).

13 Charrier, C. *et al.* Inhibition of SRGAP2 function by its human-specific paralogs induces neoteny during spine maturation. *Cell* **149**, 923-935 (2012).

14 Li, Y. *et al.* Identification of SRGAP2 as a potential oncogene and a prognostic biomarker in hepatocellular carcinoma. *Life Sci.* **277**, 119592 (2021).

15 Ji, V. & Kishore, C. The emerging roles of srGAPs in cancer. *Mol. Biol. Rep.* **49**, 755-759 (2022).

16 Liu, Y., Li, M., Yu, H. & Piao, H. lncRNA CYTOR promotes tamoxifen resistance in breast cancer cells via sponging miR-125a-5p. *Int. J. Mol. Med.* (2019).

17 Poliseno, L., Marranci, A. & Pandolfi, P. P. Pseudogenes in human cancer. *Front. Med.* **2** (2015).

18 Tian, X. *et al.* MYC-regulated pseudogene HMGA1P6 promotes ovarian cancer malignancy via augmenting the oncogenic HMGA1/2. *Cell Death Dis.* **11** (2020).

19 Qi, Y.-N., Liu, Z., Hong, L.-L., Li, P. & Ling, Z.-Q. Methyltransferase-like proteins in cancer biology and potential therapeutic targeting. *J. Hematol. Oncol.* **16** (2023).

20 Dimitrov, S. D. *et al.* Physiological modulation of endogenous BRCA1 p220 abundance suppresses DNA damage during the cell cycle. *Genes Dev.* **27**, 2274-2291 (2013).

21 Honar, Y. S. *et al.* Advanced stage, high-grade primary tumor ovarian cancer: a multi-omics dissection and biomarker prediction process. *Sci. Rep.* **13** (2023).

22 Filippou, P. S., Karagiannis, G. S. & Constantinidou, A. Midkine (MDK) growth factor: a key player in cancer progression and a promising therapeutic target. *Oncogene* **39**, 2040-2054 (2020).

23 Nash, K. T. The KISS1 metastasis suppressor: mechanistic insights and clinical utility. *Front. Biosci.* **11**, 647 (2006).

24 Jang, J.-Y., Choi, Y., Jeon, Y.-K. & Kim, C.-W. Suppression of adenine nucleotide translocase-2 by vector-based siRNA in human breast cancer cells induces apoptosis and inhibits tumor growth in vitro and in vivo. *BCR* **10**, R11 (2008).

25 Qureshi, M. A. *et al.* Pan-cancer multiomics analysis of TC2N gene suggests its important role(s) in tumourigenesis of many cancers. *APJCP* **21**, 3199-3209 (2020).

26 Li, H. *et al.* Long non-coding RNA LINC00847 induced by E2F1 accelerates non-small cell lung cancer progression through targeting miR-147a/IFITM1 axis. *Front. Med.* **8** (2021).

27 Nuñez-Olvera, S. I. *et al.* Breast cancer cells reprogram the oncogenic lncRNAs/mRNAs coexpression networks in three-dimensional microenvironment. *Cells* **11**, 3458 (2022).

28 Wang, J., Guo, W., Wang, Q., Yang, Y. & Sun, X. Recent advances of myotubularin-related (MTMR) protein family in cardiovascular diseases. *Front. Cardiovasc. Med.* **11** (2024).

29 Liang, J. *et al.* Small nucleolar RNAs: insight into their function in cancer. *Front. Oncol.* **9** (2019).

30 Luo, L. *et al.* LncRNA SNORD3A specifically sensitizes breast cancer cells to 5-FU by sponging miR-185-5p to enhance UMPS expression. *Cell Death Dis.* **11** (2020).

31 Cantor, J. R. *et al.* Physiologic medium rewires cellular metabolism and reveals uric acid as an endogenous inhibitor of UMP synthase. *Cell* **169**, 258-272 e217 (2017).

**Table S1. Composition of HPLM.** Modified from work by Cantor et. al.[31]

| Component | Concentration (µM) |
|---|---|
| Glucose | 5000 |
| Alanine | 430 |
| Arginine | 110 |
| Asparagine | 50 |
| Aspartate | 20 |
| Cysteine | 40 |
| Cystine | 100 |
| Glutamate | 80 |
| Glutamine | 550 |
| Glycine | 300 |
| Histidine | 110 |
| Isoleucine | 70 |
| Leucine | 160 |
| Lysine | 200 |
| Methionine | 30 |
| Phenylalanine | 80 |
| Proline | 200 |
| Serine | 150 |
| Threonine | 140 |
| Tryptophan | 60 |
| Tyrosine | 80 |
| Valine | 220 |
| Biotin | 0.82 |
| Choline | 21.49 |
| Folate | 2.27 |
| myo-Inositol | 194.27 |
| Niacinamide | 8.19 |
| p-Aminobenzoate | 7.29 |
| Pantothenate | 1.05 |
| Pyridoxine | 4.86 |
| Riboflavin | 0.53 |
| Thiamine | 2.96 |
| Vitamin B-12 | 0.0037 |
| 2-hydroxybutyrate | 50 |
| 3-hydroxybutyrate | 50 |
| 4-hydroxyproline | 20 |
| Acetate | 40 |
| Acetone | 60 |
| Acetylglycine | 90 |
| Alpha-aminobutyrate | 20 |
| Betaine | 70 |
| Carnitine | 40 |
| Citrate | 130 |

| | |
|---|---|
| Citrulline | 40 |
| Creatine | 40 |
| Creatinine | 75 |
| Formate | 50 |
| Fructose | 40 |
| Galactose | 60 |
| Glutathione | 25 |
| Glycerol | 120 |
| Hypoxanthine | 10 |
| Lactate | 1600 |
| Malonate | 10 |
| Ornithine | 70 |
| Pyruvate | 50 |
| Succinate | 20 |
| Taurine | 90 |
| Urea | 5000 |
| Uric acid | 350 |
| Calcium chloride | 2350 |
| Potassium chloride | 4100 |
| Magnesium chloride | 480 |
| Magnesium sulfate | 350 |
| Sodium chloride | 105000 |
| Sodium bicarbonate | 24000 |
| Disodium hydrogen phosphate | 870 |
| Calcium nitrate tetrahydrate | 40 |
| Ammonium chloride | 40 |
| Phenol red | 14 |
| Na+ | 132271 |
| K+ | 4142 |
| Ca2+ | 2390 |
| Mg2+ | 830 |
| NH4+ | 40 |
| Cl- | 116196 |
| HCO3- | 24000 |
| PO43- | 966 |
| SO42- | 350 |
| NO3- | 80 |

**Table S2. Composition of high-fat media condition.** Received from Thermo Fisher Scientific, obtained from the HPLC lipid profiles of AlbuMAX I Lipid-Rich BSA.

| Component | Concentration in high-fat media (μM) |
|---|---|
| Alpha-linolenic acid | 80 |
| Linoleic acid | 90 |
| Oleic acid | 300 |
| Stearic acid | 290 |
| Palmitic acid | 300 |

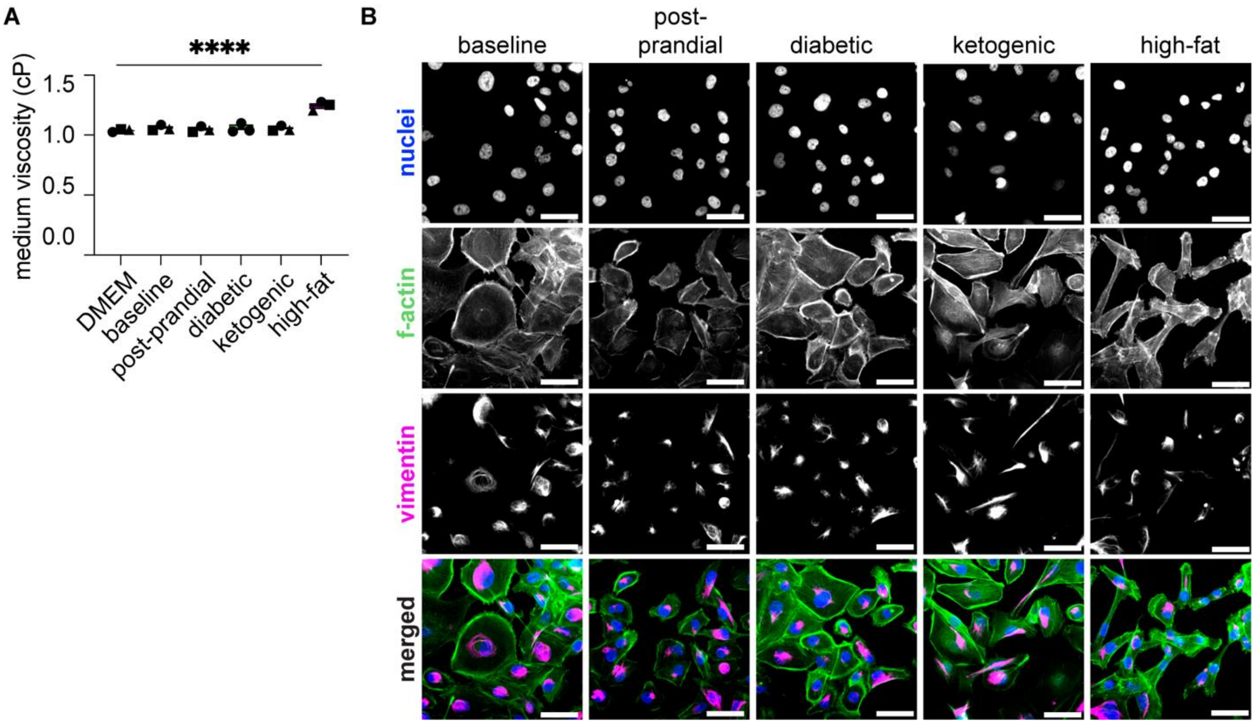


**Figure S1. Measurements of viscosity across media conditions.** (**A**) Graph showing the viscosity, in centipoise (cP), of each media formulation used in this study, pooled over n = 3 independent experiments. Each point is the average of five measurements. (**B**) Images of immunofluorescence analysis for vimentin (magenta) in cells cultured under different media conditions, counterstained for F-actin (green) and Hoechst (blue); scale bars = 50 μm. (****), p<0.0001.

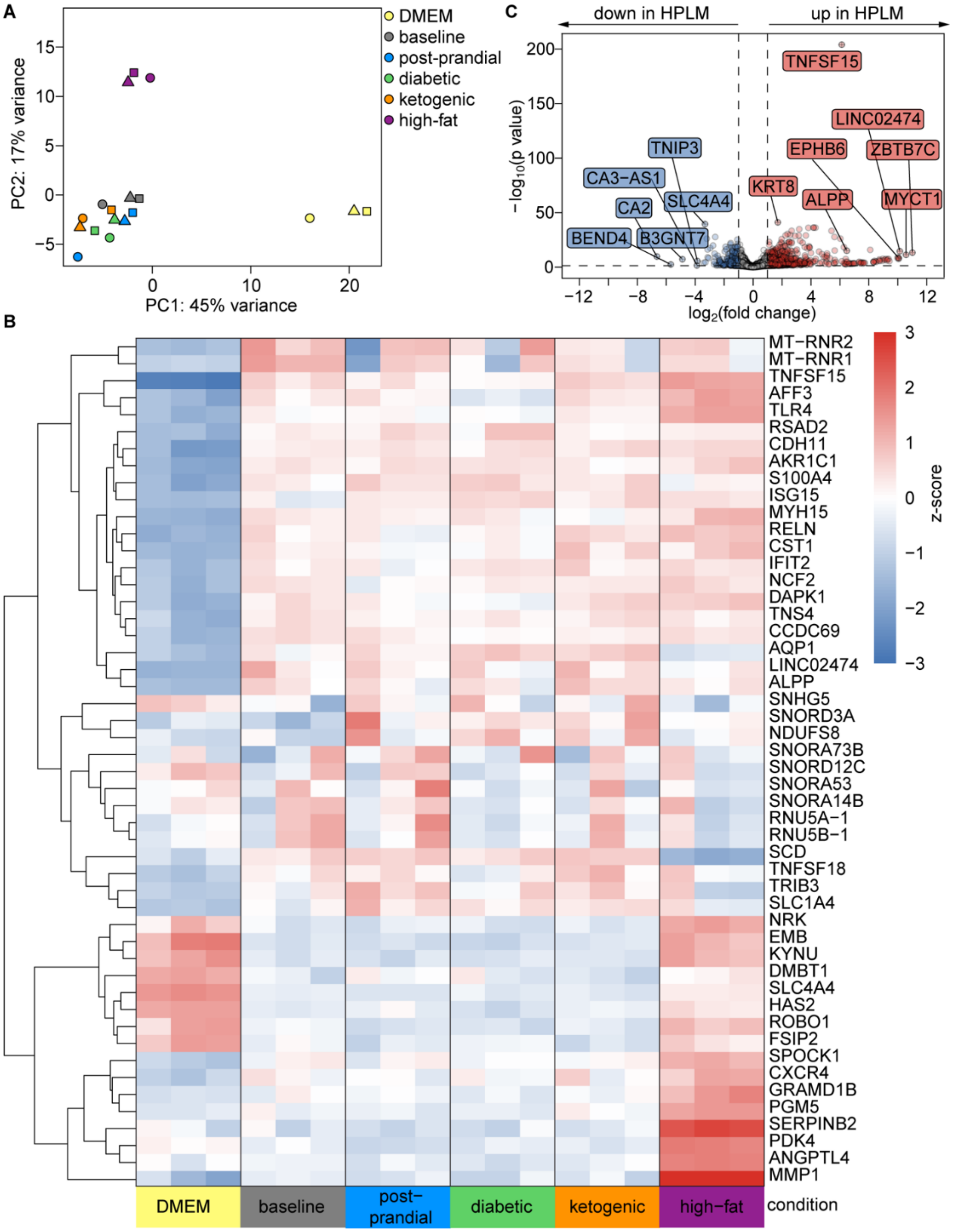


**Figure S2. Transcriptomic profiling of MDA-MB-231 cells in HPLM media conditions and standard DMEM.** (**A**) Graph of principal component analysis (PCA) based on the 1,000 most variable genes. (**B**) Heatmap displaying hierarchical clustering of differentially expressed genes across DMEM and five HPLM media conditions. Gene-expression is z-scored and color-coded by row, with red indicating higher expression and blue indicating lower expression. (**C**) Volcano plot of gene-expression changes in baseline HPLM vs. DMEM. Each point represents one gene.

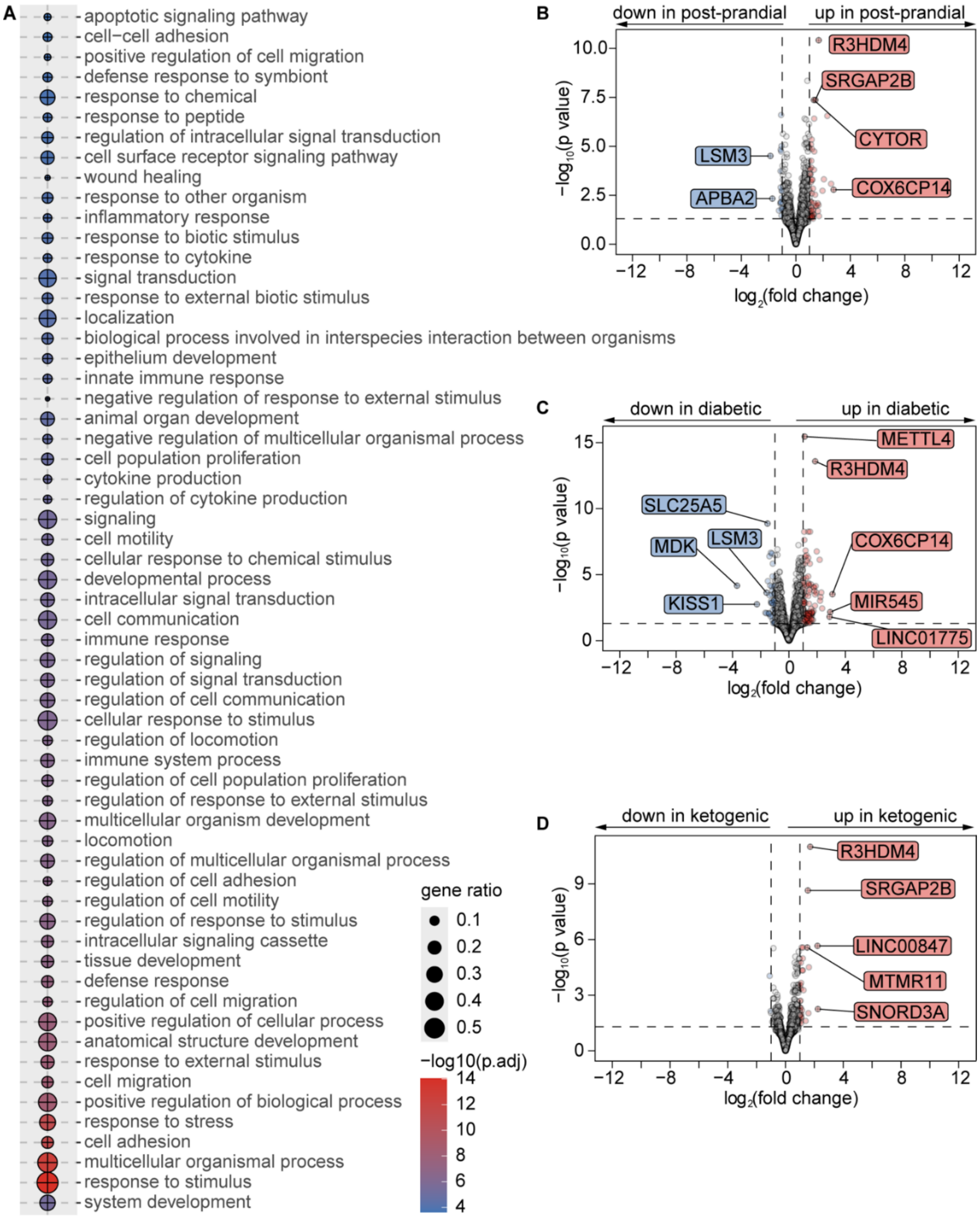


**Figure S3. Transcriptomic analysis of MDA-MB-231 cells in different media conditions.** (**A**) Over-representation analysis with the 60 most significant GO terms of differentially expressed genes in cells cultured in baseline HPLM compared to those under standard DMEM. Each dot represents a biological process, with color representing statistical significance (-log10 adjusted p-value). Upregulated (circles) processes in the baseline condition are displayed. Volcano plots comparing gene-expression between cells cultured under baseline and (**B**) post-prandial, (**C**) diabetic, and (**D**) ketogenic conditions. Genes with a fold change ≥ 2 or ≤-2 and adjusted p-value < 0.05 are highlighted in red (upregulated) and blue (downregulated) respectively.

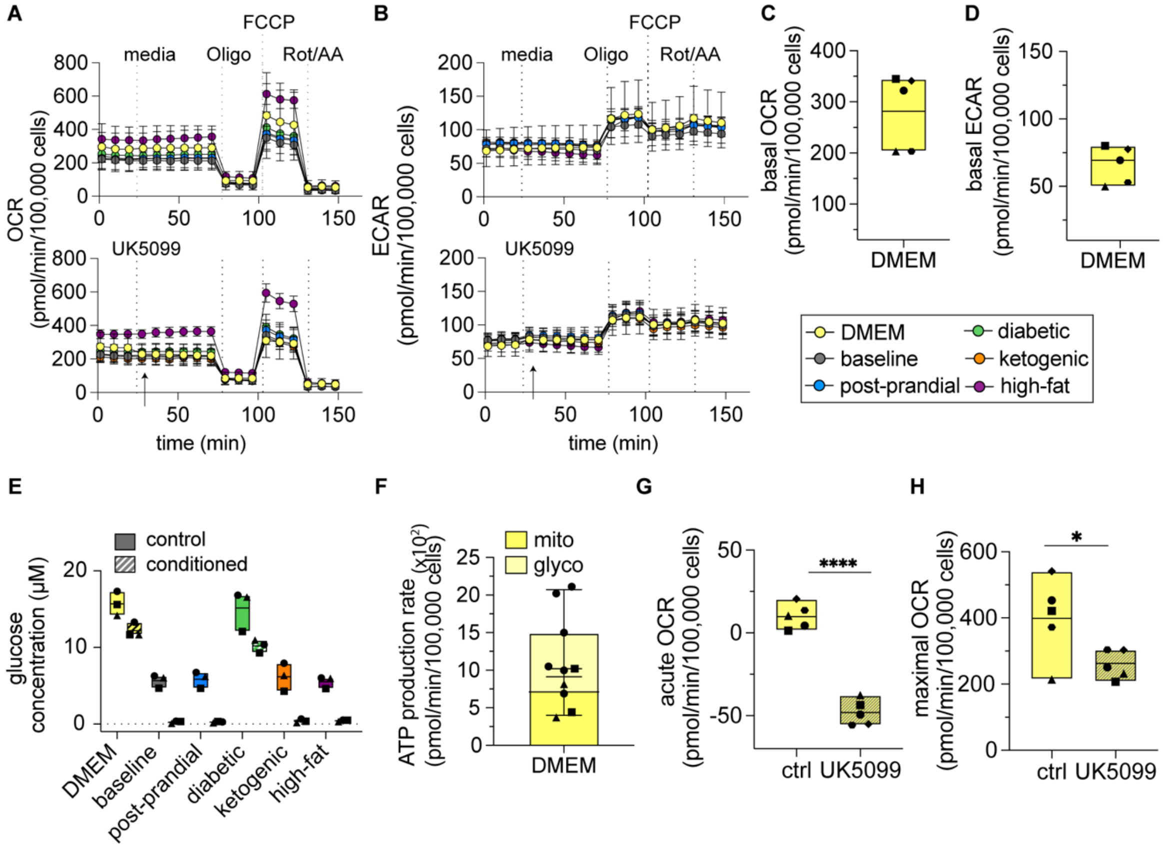


**Figure S4. Metabolic activity of MDA-MB-231 cells cultured in DMEM.** Graphs showing measurements of basal (**A**) oxygen consumption rate (OCR) and (**B**) extracellular acidification rate (ECAR) over time in cells cultured in six media conditions with or without pretreatment with UK5099. Shown are averages and standard deviations of five independent experiments. Graphs showing basal (**C**) OCR and (**D**) ECAR for cells cultured in DMEM. (**E**) Graph showing glucose levels in conditioned media after three days of culture compared to unconditioned media. Graphs showing (**F**) mitochondrial (47.6%) and glycolytic (52.4%) ATP production rates, (**G**) acute response to UK5099, and (**H**) maximal respiration after being exposed to UK5099 for cells cultured in DMEM. (*), $p<0.05$; (****), $p<0.0001$.

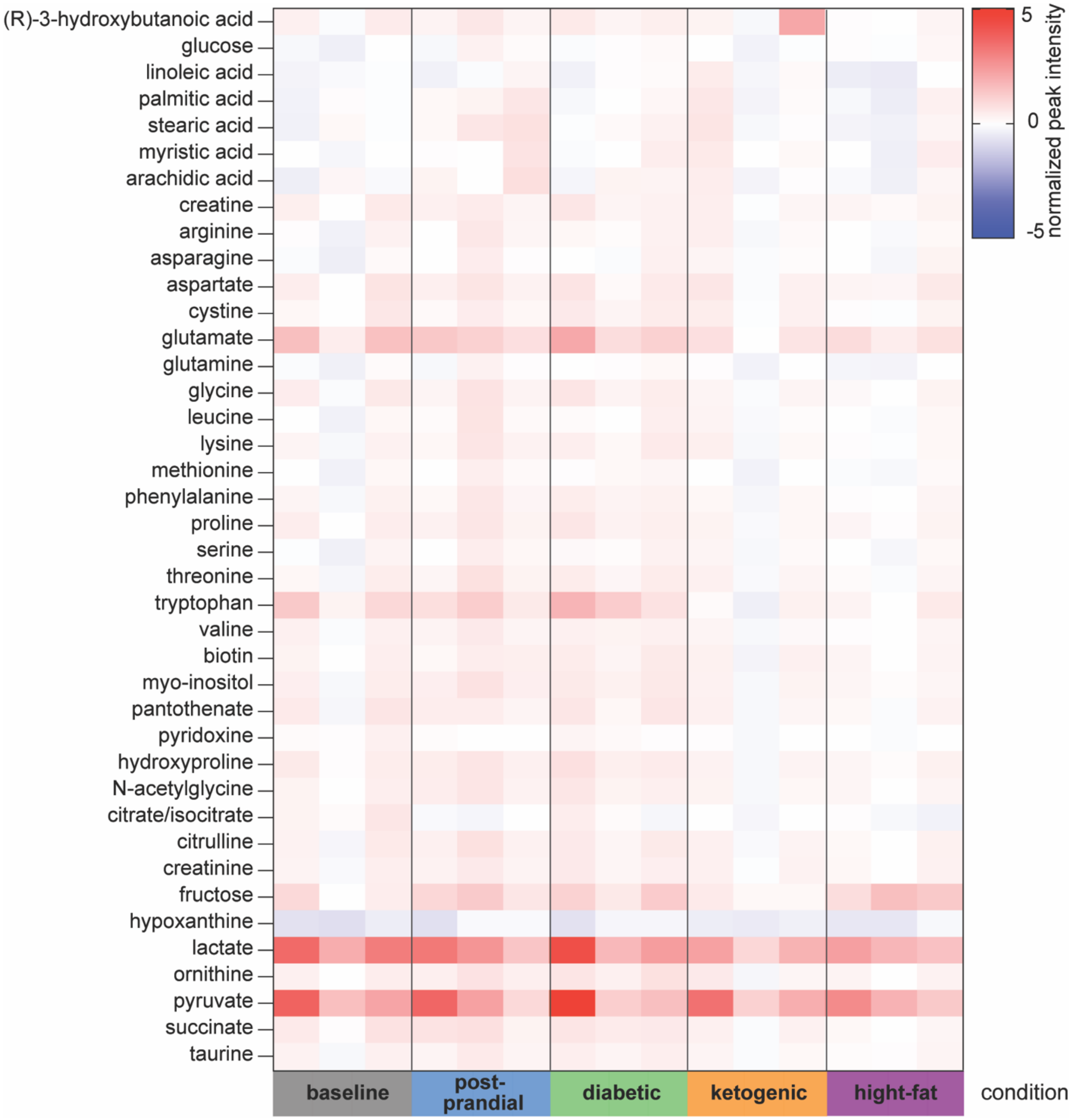


**Figure S5. Mass spectrometry analysis of conditioned media from tumors cultured under five circulating nutrient conditions.** Heatmap showing the relative abundance of metabolites in the conditioned media of engineered tumors cultured for two days under flow. Levels of metabolites were normalized to each corresponding unconditioned medium (see Methods).

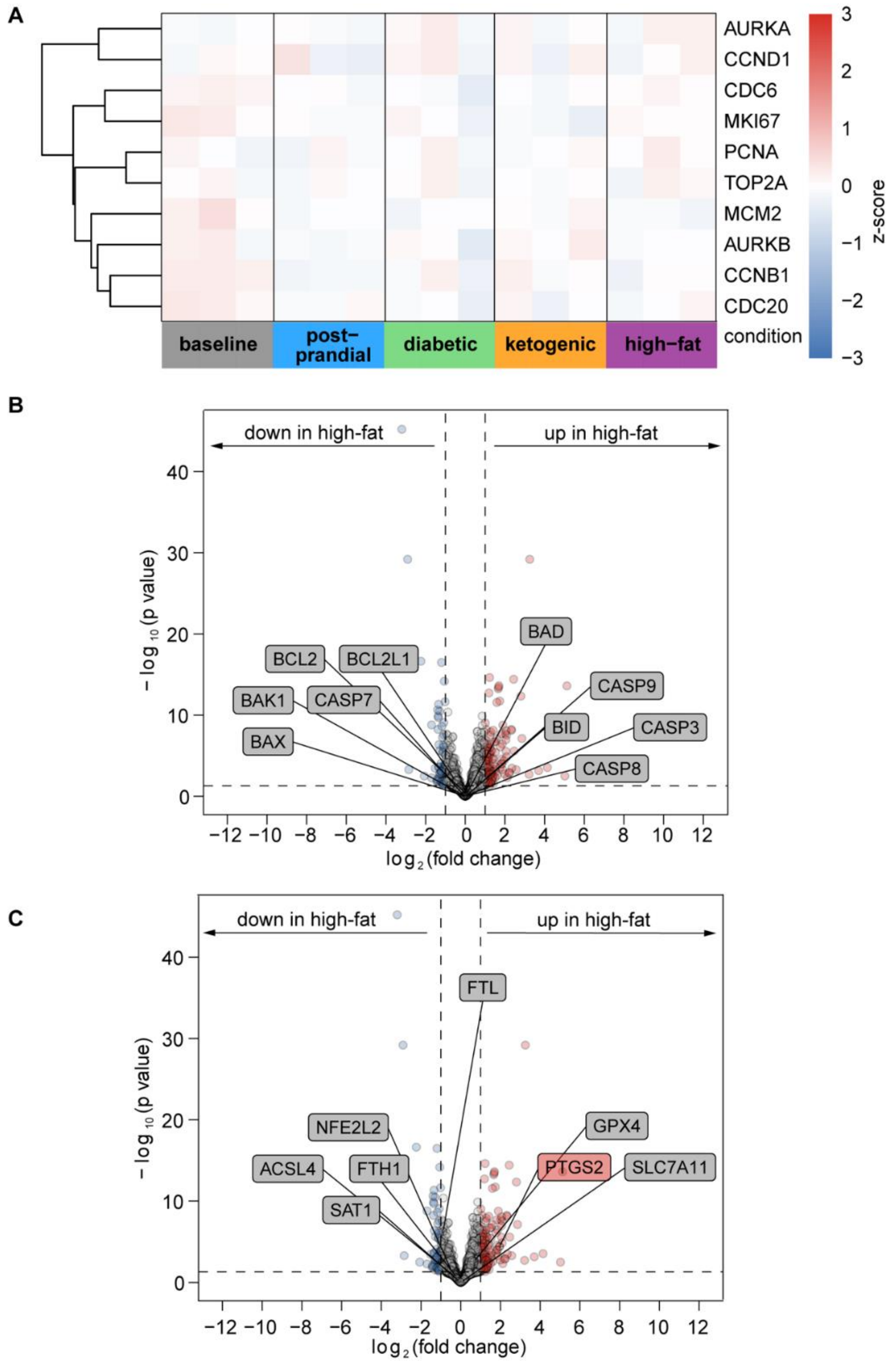


**Figure S6. Gene-expression analysis reveals no significant differences in markers of proliferation, apoptosis, or ferroptosis across media mimicking circulating nutrient conditions.** (**A**) Heatmap showing the expression of key proliferation-associated genes across five media conditions. Volcano plots comparing gene-expression between high-fat and baseline conditions, highlighting genes related to (**B**) apoptosis and (**C**) ferroptosis. Most highlighted genes fall below the ±2-fold change region of the volcano plot, indicating no significant differences in their expression levels between the conditions.

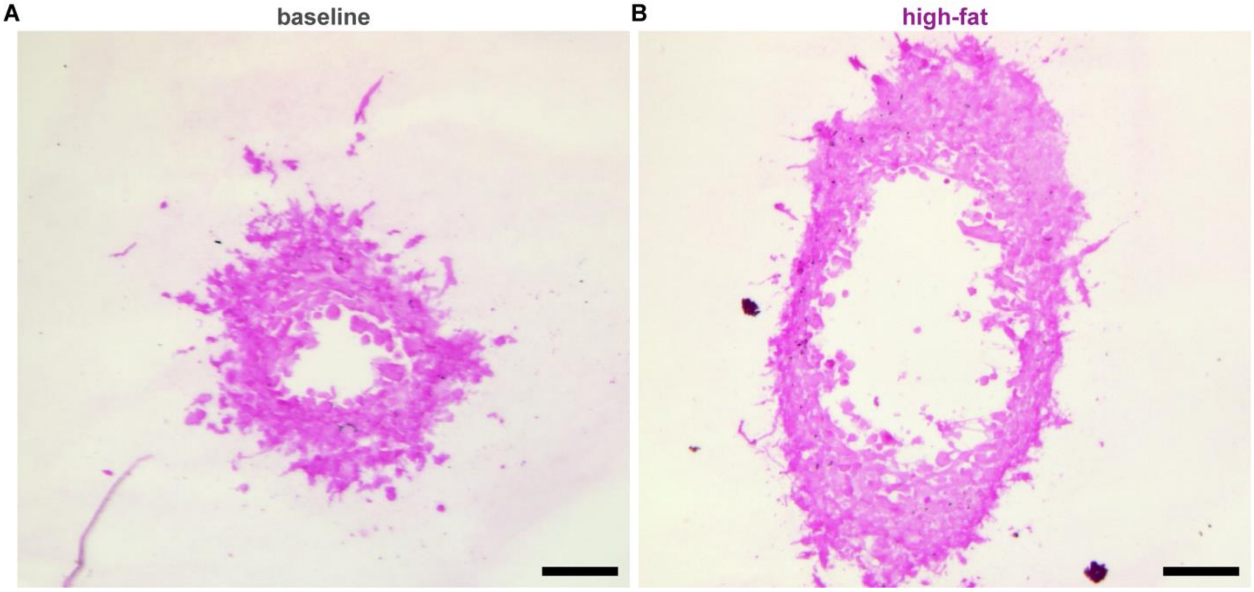


**Figure S7. Prussian blue staining reveals no detectable iron in baseline or high-fat tumor sections.** Images of Prussian blue staining of nuclei (pink) and iron (blue) in sections of (**A**) baseline and (**B**) high-fat tumor sections at day six of culture; scale bars = 100 µm.
